# Proteolysis of fibrillin-2 microfibrils is essential for normal skeletal development

**DOI:** 10.1101/2021.02.03.429587

**Authors:** Timothy J. Mead, Daniel R. Martin, Lauren W. Wang, Stuart A. Cain, Cagri Gulec, Elisabeth Cahill, Joseph Mauch, Dieter P. Reinhardt, Cecilia W. Lo, Clair Baldock, Suneel S. Apte

## Abstract

The embryonic extracellular matrix (ECM) undergoes transition to mature ECM as development progresses, yet specific transition mechanisms ensuring ECM proteostasis and their regulatory impact are poorly defined. Fibrillin microfibrils are macromolecular ECM complexes serving structural and regulatory roles. In mice, *Fbn1* and *Fbn2,* encoding major microfibrillar components, are strongly expressed during embryogenesis, but fibrillin-1 is the major component observed in adult tissue microfibrils. Here, analysis of mouse *Adamts6* and *Adamts10* mutant embryos, lacking these homologous secreted metalloproteases individually and in combination, along with in vitro analysis of microfibrils, measurement of ADAMTS6-fibrillin affinities and N-terminomics identification of ADAMTS6-cleaved sites, demonstrates a transcriptionally adapted system for fibrillin-2 proteolysis that contributes to postnatal fibrillin-1 dominance. The lack of ADAMTS6, alone and in combination with ADAMTS10 led to excess fibrillin-2 in perichondrium, with impaired skeletal development resulting from a drastic reduction of aggrecan, cartilage link protein and impaired BMP, but not TGFβ signaling in cartilage. Although ADAMTS6 cleaves fibrillin-1 and fibrillin-2 as well as fibronectin, which provides the initial scaffold for microfibril assembly, primacy of the protease-substrate relationship between ADAMTS6 and fibrillin-2 was unequivocally established by reversal of these defects in *Adamts6*^-/-^ embryos by genetic reduction of *Fbn2*, but not *Fbn1*.

## Introduction

In addition to proliferation and differentiation of resident cells, proper tissue and organ development, structure and function require an appropriate extracellular matrix (ECM). How ECM architecture and stoichiometry are maintained and adjusted coordinately with the dynamic nature of morphogenesis and transition into the adult organism is unknown. Ontogenetically, the earliest cell collectives formed sheets and tubes with a well-established basement membrane that provided a substrate for cell migration and adhesive inputs that determined cell polarity. Expansion of ECM-encoding genes enabled formation of a complex interstitial matrix that promoted evolution of ever more complex organisms (1), but presented a challenge for remodeling of the increasingly diverse ECM repertoire within complex ECM assemblies. One solution may have been the concomitant expansion of genes encoding secreted and cell-surface proteases (2, 3). For example, of 19 ADAMTS proteases in mammals compared to only 2 in the fruitfly, the majority cleave ECM molecules (4, 5). However, which specific ECM molecules and supramolecular assemblies are targets of the individual proteases, the specific contexts they operate in, and possible cooperative/regulatory relationships between individual proteases are poorly understood. The embryonic interstitial ECM is highly hydrated owing to abundant macromolecular hyaluronan (HA)-proteoglycan aggregates whereas fibrillar components, primarily collagens and elastin, dominate juvenile and adult ECM composition to provide structural resilience compatible with the greater mechanical demands imposed during postnatal life. Beyond structural roles, ECM sequesters and regulates the activity of morphogens and growth factors (6, 7), and ECM proteolysis can generate bioactive fragments, termed matrikines (8).

Fibrillin microfibrils have a crucial role in tissue development and homeostasis by providing mechanical stability and limited elasticity to tissues and/or regulating growth factors of the TGFβ superfamily, including bone morphogenetic proteins (BMPs) and growth/differentiation factors (GDFs) (6, 7), along with a key role in elastic fiber assembly (9, 10). Fibrillins are large, cysteine-rich glycoproteins containing many epidermal growth factor (EGF)-like repeats. Of the three fibrillin isoforms, fibrillin-2 and fibrillin-3 (in humans) are primarily expressed during embryogenesis (11–13). The gene encoding fibrillin-3 is inactivated in mice (14), providing a simpler scenario than in humans for investigating developmental regulation of microfibril composition and the role of proteolytic turnover therein. Among numerous gene mutations affecting the skeleton (15), *FBN1* and *FBN2* mutations cause distinct dominantly-inherited human connective tissue disorders, Marfan syndrome and congenital contractural arachnodactyly, respectively (16). Despite overlapping features such as skeletal overgrowth and poor muscular development, each disorder mainly has distinct manifestations, indicating that fibrillin isoforms may contribute specific properties to microfibrils, have a tissue-specific function, or form distinct ECM networks. Severe cardiovascular manifestations, especially potentially lethal aortic root and ascending aorta aneurysms as well as ocular manifestations, occur frequently in Marfan syndrome, but neither is typically associated with *FBN2* mutations (16). In mice, *Fbn2* deficiency affects myogenesis and distal limb patterning, reflecting a role for fibrillin-2 in BMP regulation (17, 18). Fibrillin microfibrils can be homotypic or heterotypic assemblies of fibrillin-1 and/or fibrillin-2 (19–21), but since each fibrillin appears to have distinct roles in vivo as well as in vitro (22), an intriguing question is how their correct stoichiometry is maintained, and what impact an excess of fibrillin microfibrils or altered fibrillin isoform stoichiometry may have on biological systems. Here, analysis of mouse mutants of the homologous secreted metalloproteases ADAMTS6 and ADAMTS10 has uncovered a transcriptionally adapted system for ECM proteostasis and defined functional specificity of the ADAMTS6-fibrillin relationships.

ADAMTS10 has established relevance to skeletal dysplasias and Marfan syndrome. Recessive *ADAMTS10* mutations lead to an acromelic dysplasia, Weil-Marchesani syndrome 1 (WMS1) (23), whereas dominant *FBN1* mutations cause a similar disorder, WMS2 (24), strongly suggesting a functional relationship between ADAMTS10 and fibrillin-1 (25, 26). ADAMTS10 bound fibrillin-1 directly and accelerated fibrillin-1 microfibril biogenesis in vitro, cleaving fibrillin-inefficiently (27, 28). Mice with targeted *Adamts10* inactivation or homozygous for a human WMS-causing *ADAMTS10* mutation had impaired long bone growth and increased muscle mass along with reduced fibrillin-1 staining in skeletal muscle and persistence of fibrillin-2 microfibrils in skeletal muscle and the eye (29, 30). ADAMTS10 undergoes inefficient processing by furin, typically a prerequisite for activation of ADAMTS proteases. However when the ADAMTS10 furin processing site was experimentally optimized to favor propeptide cleavage, it proteolytically processed fibrillin-2 (30). In contrast, the ADAMTS10 homolog, ADAMTS6, is efficiently processed by furin (27, 31), but its activity and functional relationship with ADAMTS10 are poorly characterized, although it is known to cleave latent TGFβ-binding protein 1, which is a fibrillin relative, as well as cell-surface heparan sulfate proteoglycans (27). Since ADAMTS proteases cooperate in several physiological contexts where they are co-expressed (32–37) and their pairwise homology hsuggested the possibility of transcriptional adaptation (38, 39), which has not been systematically addressed in this family, we investigated the impact of single or combined *Adamts6* and *Adamts10* inactivation on mouse skeletal development, and defined the underlying mechanisms. The findings provide intriguing insights into fibrillin microfibril proteostasis.

## Results

### Transcriptional adaptation of *Adamts6* and *Adamts10*

Previous work had suggested that knockdown of *ADAMTS10* in cultured human ARPE-19A cells increased *ADAMTS6* expression (27). To investigate the possibility that germline inactivation of either mouse gene affected expression of the other, *Adamts6* and *Adamts10* mRNA levels were measured in limbs, heart and lungs of *Adamts6*^-/-^ and *Adamts10*^-/-^ mice. qRT-PCR analysis showed that *Adamts6* mRNA was not altered with respect to wild type levels in *Adamts6-*mutant mice (which have a missense Ser^149^Arg mutation that abrogates protein secretion, essentially comprising a knockout (31)), but was consistently increased in *Adamts10*^-/-^ tissues (Figure 1A). *Adamts10* mRNA was significantly reduced in *Adamts10*^-/-^ mice (which have an intragenic IRES-*lacZ* insertion that disrupts mRNA continuity), but was unaltered in *Adamts6*^-/-^ tissues (Figure 1A). In addition to the transcriptional adaption of this gene pair, we asked whether ADAMTS6 and ADAMTS10 cleaved the other. Neither ADAMTS10 nor furin-optimized ADAMTS10 cleaved ADAMTS6 and ADAMTS6 did not cleave ADAMTS10 (Figure 1 – figure supplement 1). These finding raised two possibilities, i) That *Adamts10*^-/-^ phenotypes could have been buffered by a compensating increase in *Adamts6* mRNA and activity, and ii) That cooperative functions, such as were previously identified in other combined mutants of homologous ADAMTS proteases (35, 36), may exist. Both possibilities could be addressed by generation of *Adamts6*^-/-^;*Adamts10*^-/-^ mice. Therefore, after initial characterization of skeletal development in *Adamts6* mutant mice, we analyzed the combined null mutants. Each mutant genotype was recovered at the expected Mendelian ratio at the end of the embryonic period (Figure 1 – figure supplement 2).

**Figure 1.**
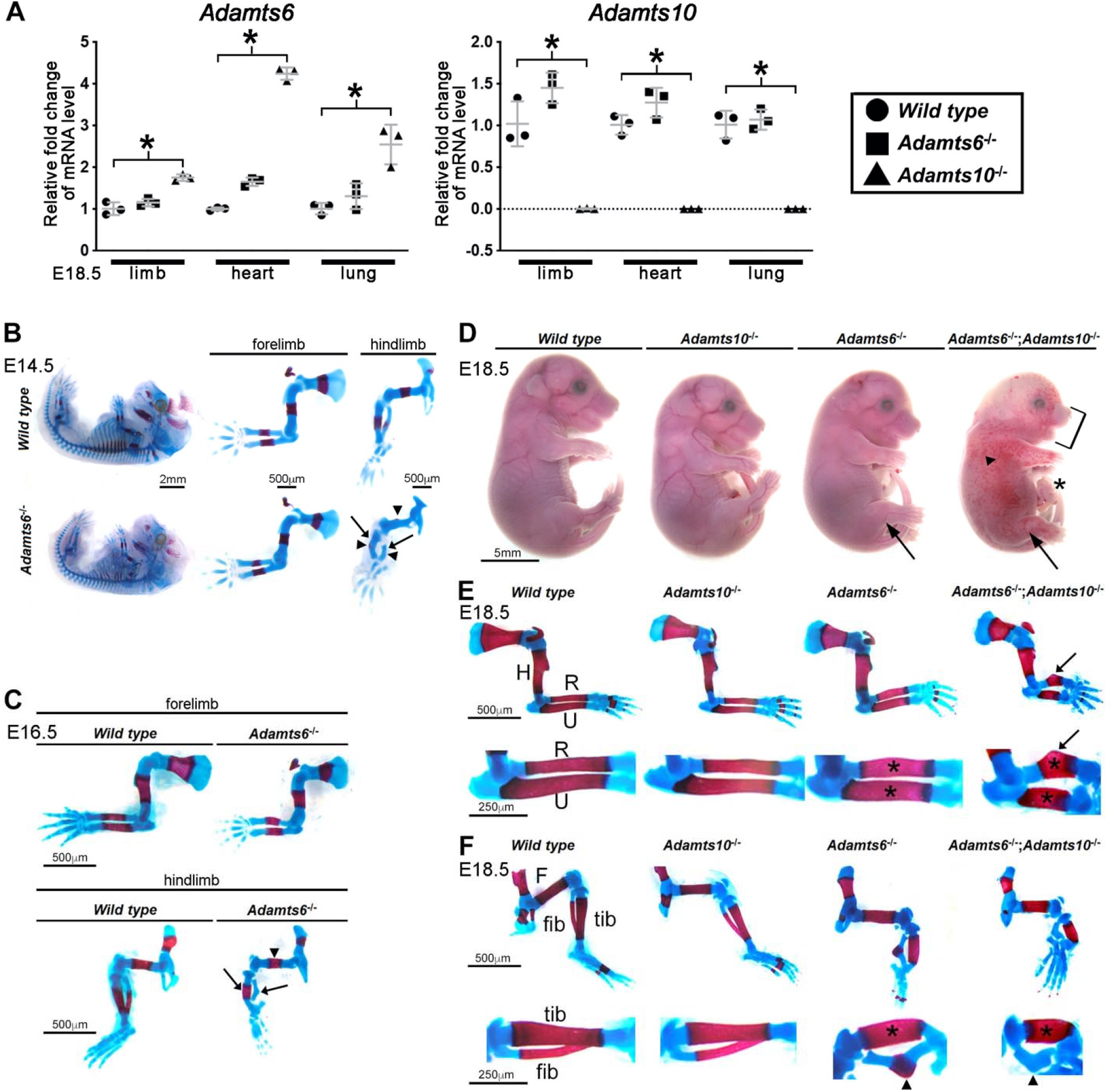
ADAMTS6 and ADAMTS10 are subject to transcriptional adaptation and cooperate in skeletal development. **(A)** qRT-PCR analysis of *Adamts6* and *Adamts10* mRNA levels in wild type, *Adamts6^-^*^/-^ and *Adamts10*^-/-^ limb, heart and lung show that *Adamts6* mRNA is elevated in *Adamts10*^-/-^ tissues, whereas *Adamts10* mRNA is not significantly altered in *Adamts6^-^*^/-^ tissues (n=3). Error bars represent ± SEM. *p ≤0.01, Student *t* test). **(B, C)** E14.5 **(B)** and E16.5 **(C)** alcian blue- and alizarin red-stained skeletons show severely short and under-ossified *Adamts6^-^*^/-^ hindlimb skeleton (arrowheads) with distorted tibia and fibula (arrow). *Adamts6^-^*^/-^ forelimbs are not as severely affected as hindlimbs. **(D)** E18.5 *Adamts6^-^*^/-^;*Adamts10*^-/-^ embryos have more severe hindlimb malformations (arrows), shorter forelimbs, short snout and mandibular hypoplasia (bracket) an omphalocele (asterisk) and widespread cutaneous hemorrhage (arrowhead) compared to *Adamts6^-^*^/-^ and *Adamts10*^-/-^ embryos. **(E, F)** Alcian blue- and alizarin red-stained *Adamts6^-^*^/-^;*Adamts10*^-/-^ forelimbs **(E)** and hindlimbs **(F)** show unremodeled, thickened (asterisks) and shortened long bones (H, humerus; F, femur; U, ulna; R, radius; Tib, tibia; Fib, fibula) with a bent radius (arrow) and tibia (arrowhead). Note the unossified fibula in the *Adamts6^-^*^/-^;*Adamts10*^-/-^

### Severe skeletal malformations in *Adamts6-*deficient mice are exacerbated by combined *Adamts10* inactivation

*Adamts6*^-/-^ embryos had severe reduction of crown to rump length, which was also statistically significant in *Adamts10*^-/-^ mice (Figure 1 – figure supplement 3A). Whereas *Adamts6*^-/-^ forelimbs appeared shorter, *Adamts6*^-/-^ hindlimbs were severely short and internally rotated (Figure 1B, C, D, F, Figure 1 – figure supplement 3B). Alizarin red- and alcian blue-stained skeletal preparations at embryonic day (E)14.5, when ossification is initiated, and at E16.5 and E18.5, by which time it is established, showed reduced ossification of *Adamts6*^-/-^ hindlimb distal long bones and skeletal deformation, (Figure 1B, C, F), most evident in the thicker tibia and angulated fibula (Figure 1B, C, F, arrows). *Adamts6*^-/-^ forelimbs demonstrated failure of diaphyseal remodeling (e.g., wider, tubular-appearing radius and ulna) and smaller ossific centers than wild type littermates (Figure 1E). Neither forelimbs nor hindlimbs showed skeletal patterning anomalies. In contrast to *Adamts6*^-/-^ embryos, *Adamts10*^-/-^ embryos had mild shortening of individual long bones, which was not statistically significant (29, 30). The axial and craniofacial skeleton were also abnormal in E18.5 *Adamts6*^-/-^ mice, with shortened, tubular ribs, lack of sternal segmentation and an under-ossified xiphoid process (Figure 1 – figure supplement 4). Additionally, their vertebral bodies were smaller in size with corresponding size reduction of all vertebral ossification centers. *Adamts6*^-/-^ craniofacial skeletons had reduced anterior-posterior and nasal-occipital dimensions, corresponding reduction in size of individual elements, delayed ossification of parietal and other bones and wider anterior fontanelles (Figure 1 – figure supplement 4).

Histologic comparison of alcian blue-stained E14.5 wild type and *Adamts6*^-/-^ long bone sections showed delayed endochondral ossification, with persistence of hypertrophic chondrocytes and lack of vascular invasion in the primary ossification centers (Figure 1 – figure supplement 5A). E18.5 alcian blue-stained sections revealed under-ossified and malformed *Adamts6*^-/-^ tibia and fibula (Figure 1 – figure supplement 5B). *Adamts6*-deficient E18.5 distal femoral and proximal tibial cartilage had a histologically expanded hypertrophic chondrocyte zone, presumably reflecting delayed ossification within primary centers (Figure 1 – figure supplement 5C).

To determine possible cooperative roles of ADAMTS6 and ADAMTS10, mice with combinations of the two mutant alleles were obtained by interbreeding, since each gene is on a distinct mouse chromosome. *Adamts6*^-/-^;*Adamts10*^-/-^ embryos demonstrated markedly more severe anomalies than *Adamts6*^-/-^ mutants including subcutaneous hemorrhage, micrognathia and an omphalocele, along with more severe forelimb and hindlimb dysmorphology (Figure 1D). Skeletal preparations showed more severe hindlimb anomalies than in *Adamts6^-/-^* mutants and appearance of forelimb abnormalities similar in degree of severity to *Adamts6^-/-^* hindlimb mutants, with externally evident shortening, the greatest reduction of nose-rump length among all genotypes and the shortest long bones among the generated genotypes (Figure 1D-F, Figure 1 – figure supplement 2, Figure 1 - figure supplement 3B, Figure 1 - figure supplement 5B-C).

Fibular ossification was minimal, and the zeugopod was angulated with pronounced tibio-fibular bending (Figure 1F). The shortened and tubular ribs, vertebral bodies with smaller ossification centers, lack of xiphoid process ossification and poor ossification of cranial bones resulting in larger fontanelles, further demonstrated more severe skeletal malformations than *Adamts6*^-/-^ embryos (Figure 1 – figure supplement 4). Whereas inactivation of one *Adamts6* allele in *Adamts10*^-/-^ mice did not significantly worsen the observed dysmorphology and skeletal phenotype, *Adamts10* haploinsufficiency further reduced *Adamts6*^-/-^ crown to rump length and led to shorter long bones (Figure 1 – figure supplement 3B). Thus, both transcriptional adaptation and cooperativity of ADAMTS6 and ADAMTS10 appear to have a role in ensuring normal skeletal development.

### *Adamts6* and *Adamts10* have overlapping expression in limb skeletal development

To follow up on prior publications showing strong expression of *Adamts6* in the heart and of *Adamts10* in multiple embryonic and adult tissues (30, 31, 40) as well as immunohistochemical localization of ADAMTS10 in limb growth cartilage, perichondrium and muscle (41), we compared the spatial and temporal localization of their mRNAs during hindlimb development using RNAScope in situ hybridization (*Adamts6*) and an intragenic lacZ reporter (*Adamts10*). *Adamts6* and *Adamts10* exhibited overlapping expression in resting and proliferating zone chondrocytes and perichondrium of E13.5 and E14.5 hindlimb long bones (Figure 1 – figure supplement 6). At E14.5, when the first primary ossific centers are formed, both genes were expressed at sites of vascular invasion of cartilage (Figure 1 – figure supplement 6B, arrowheads). *Adamts6* expression was also noted in tendons and skeletal muscle around the knee joints and *Adamts10* was ubiquitously expressed throughout the joint interzone (Figure 1 – figure supplement 6).

### *Adamts6*-deficient skeletal elements have disorganized growth plate cartilage and dramatically reduced proteoglycan

RGB trichrome-stained E18.5 sections revealed disorganized *Adamts6*^-/-^ and *Adamts6*^-/-^; *Adamts10*^-/-^ growth plate chondrocytes with a dramatic reduction in staining with the alcian blue component, which detects sulfated proteoglycans (Figure 2A). *Adamts6*^-/-^ and *Adamts6*^-/-^;*Adamts10*^-/-^ resting zone (RZ) chondrocytes were tightly packed compared to wild type with disorganized columnar zone (CZ) and hypertrophic zone (HZ) chondrocytes, whereas *Adamts10*^-/-^ growth plate chondrocytes resembled the wild type. To define cartilage proteoglycan content further, sections were stained with aggrecan and cartiage link protein antibodies, which revealed less intense staining in *Adamts6*^-/-^ and *Adamts6*^-/-^;*Adamts10*^-/-^ cartilage as compared to control and *Adamts10^-/-^*; this was especially evident in the RZ (Figure 2B). Consistent with the lack of aggrecan,we observed reduced staining of the transcription factor Sox9, whose expression drives chondrogenesis. In contrast, the HZ marker collagen X revealed comparable matrix staining in all genotypes, and suggested increased HZ thickness in *Adamts10*^-/-^, *Adamts6*^-/-^ and *Adamts6*^-/-^;*Adamts10*^-/-^ femur, potentially related to delayed ossification (Figure 2B, Figure 1 – figure supplement 5C). PCNA and TUNEL staining revealed no change in cell proliferation or cell death, respectively, in *Adamts6*^-/-^ femoral cartilage (Figure 2 – figure supplement 1).

**Figure 2:**
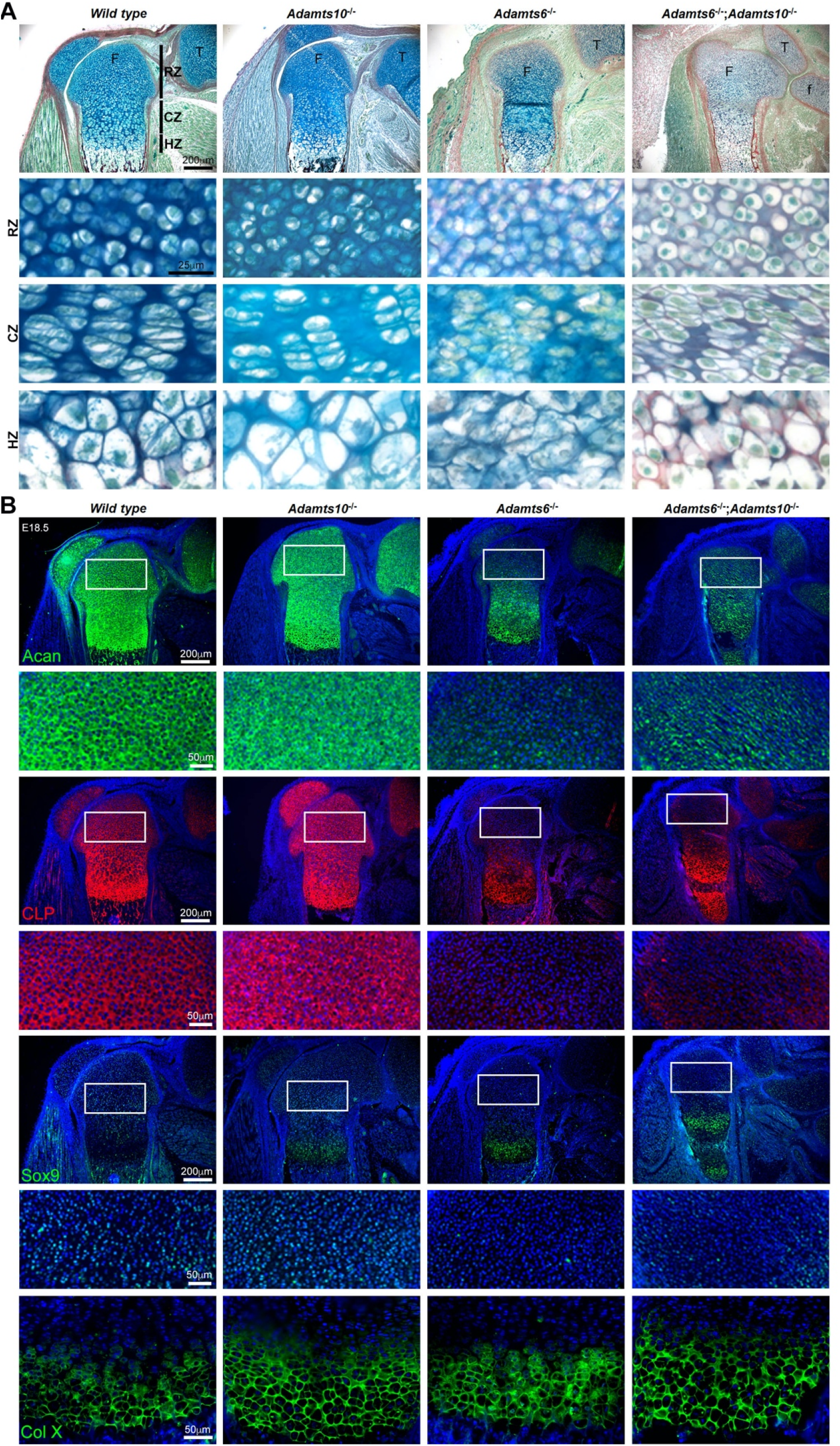
*Adamts6*-deficent hindlimbs have disorganized chondrocytes and reduced cartilage proteoglycan. **(A)** RGB trichrome-stained E18.5 knees show disorganized chondrocytes and reduced alcian blue staining in *Adamts6*^-/-^ and *Adamts6^-/^*^-^;*Adamts10*^-/-^ bones. F, femur; T, tibia; f, fibula; RZ, resting zone; CZ, columnar zone; HZ, hypertrophic zone. **(B)** Severely reduced aggrecan (Acan), cartilage link protein (CLP) and Sox9 in *Adamts6^-/^*^-^ and *Adamts6^-/^*^-^;*Adamts10*^-/-^ cartilage and wider collagen X stained zone (Col X) indicating an expanded HZ.

### Fibrillin*-2* accumulates in *Adamts6*^-/-^*, Adamts10*^-/-^ and *Adamts6*^-/-^;*Adamts10*^-/-^ hindlimbs

Since prior work had shown that ADAMTS10 had a strong functional relationship with fibrillin microfibrils, specifically acceleration of fibrillin-1 assembly and fibrillin-2 proteolysis (albeit by furin site-optimized ADAMTS10) (28–30), we investigated changes in fibrillin-1/-2 distribution and staining intensity using immunofluorescence. First, we used microfibril-associated glycoprotein 1 (MAGP1), which binds to both fibrillin-1 and fibrillin-2 (42), to report their combined abundance and detected increased staining intensity in in *Adamts10*^-/-^, *Adamts6*^-/-^ and *Adamts6*^-/-^;*Adamts10*^-/-^ femur perichondrium (Figure 3A). Immunostaining with monospecific fibrillin antibodies showed more intense fibrillin-2 staining in *Adamts10*^-/-^, *Adamts6*^-/-^ and *Adamts6*^-/-^;*Adamts10*^-/-^ perichondrium (Figure 3B) but no consistent difference in fibrillin-1 staining (Figure 3C). *Fbn1* and *Fbn2* mRNA levels were unchanged in *Adamts6*- and *Adamts10*-deficient hindlimbs, suggesting that increased fibrillin-2 staining was not a result of increased transcription (Figure 3 – figure supplement 1).

**Figure 3:**
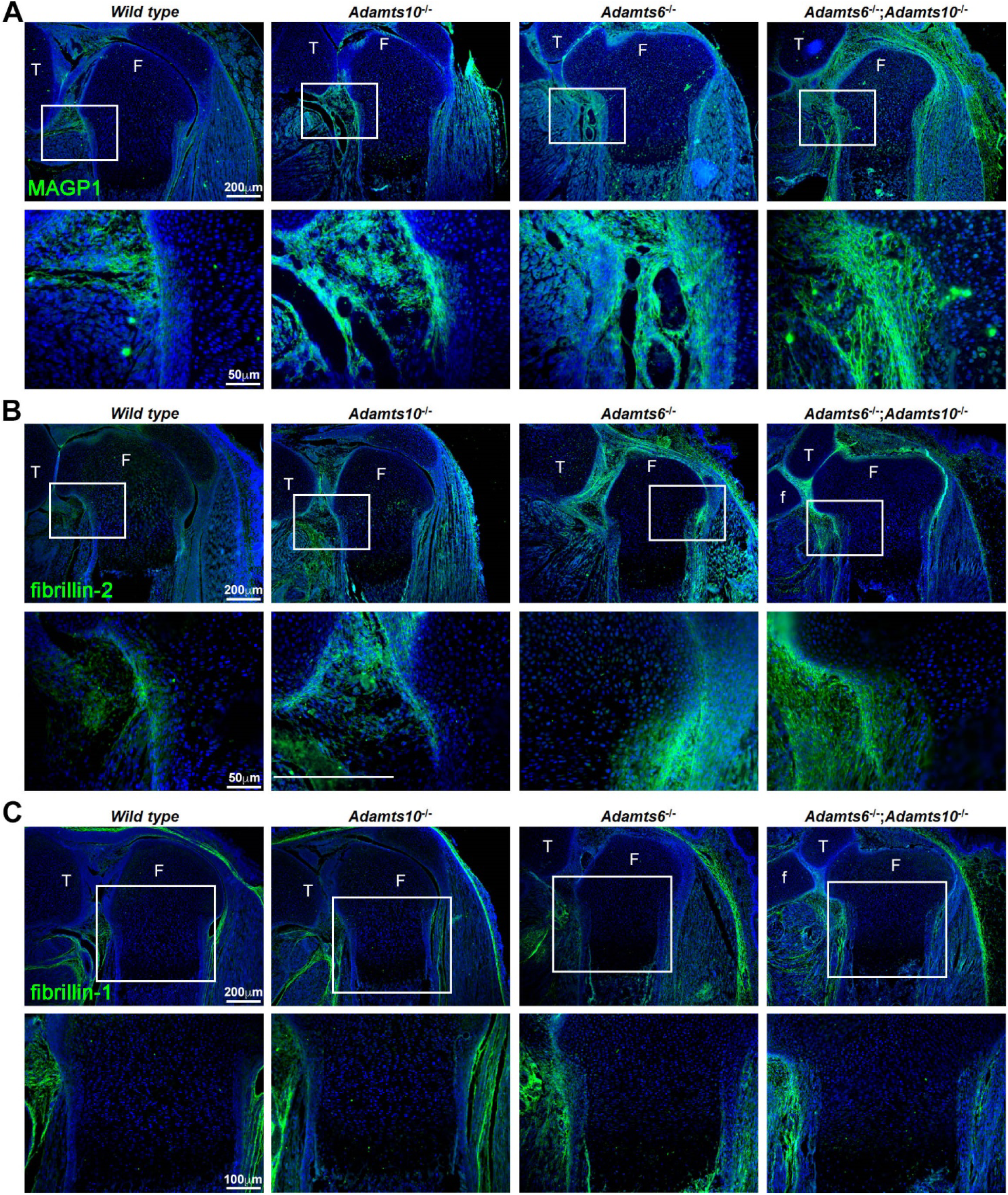
Increased MAGP1 and fibrillin-2 staining in *Adamts6^-/-^;Adamts10^-/-^* perichondrium. **(A-B)**Increased staining intensity (green) of MAGP1 (**A**) and fibrillin-2 (**B**) in E18.5 *Adamts6*- and *Adamts10*- deficient knee joints. **(C)** No consistent change in fibrillin-1 staining (green) was seen between the various genotypes. Images are representative of n=3. Sections are counterstained with DAPI (blue). F, femur; T, tibia; f, fibula.

### ADAMTS6 binds to fibrillin-2 microfibrils formed by cultured fibroblasts

We investigated ADAMTS6 binding to homotypic fibrillin-2 microfibrils assembed by *Fbn1*^-/-^ mouse embryo fibroblasts (MEFs). Co-cultures of *Fbn1*^-/-^ MEFs with HEK293T cells stably expressing catalytically inactive ADAMTS6 (ADAMTS6 Glu^404^Ala, referred to hereafter as ADAMTS6 EA) for 6 days illustrated specific co-localization of ADAMTS6 EA with fibrillin-2 microfibrils (Figure 4, top rows). When *Fbn1*^-/-^ MEFs were co-cultured with HEK293F cells expressing active ADAMTS6 (2nd row), no fibrillin-2 staining was seen, suggestive of proteolytic degradation of fibrillin-2 microfibrils or interference with their assembly. Similarly, 4 day-old cultures of *Fbn2*^-/-^ mouse skin fibroblasts (MSF) revealed co-localization of ADAMTS6 EA with fibrillin-1 and demonstrated absence of fibrillin-1 microfibrils in the presence of ADAMTS6 (Figure 4, middle panels). Since a fibronectin fibrillar matrix is formed soon after plating of cells and acts as a template for fibrillin-microfibril assembly (43, 44), we similarly analyzed fibronectin staining, observing that ADAMTS6 EA co-localized with fibronectin and that fibronectin fibrils were absent in the presence of ADAMTS6 (Figure 4, bottom panels). Furthermore, MEFs isolated from *Adamts6*^-/-^ embryos showed greater fibrillin-2, fibrillin-1 and fibronectin microfibril staining intensity than wild type MEFs (Figure 4 – figure supplement 1). Together, these findings suggested that ADAMTS6 may bind to and proteolytically degrade fibronectin and fibrillin microfibrils or unassembled fibrillin and fibronectin.

**Figure 4:**
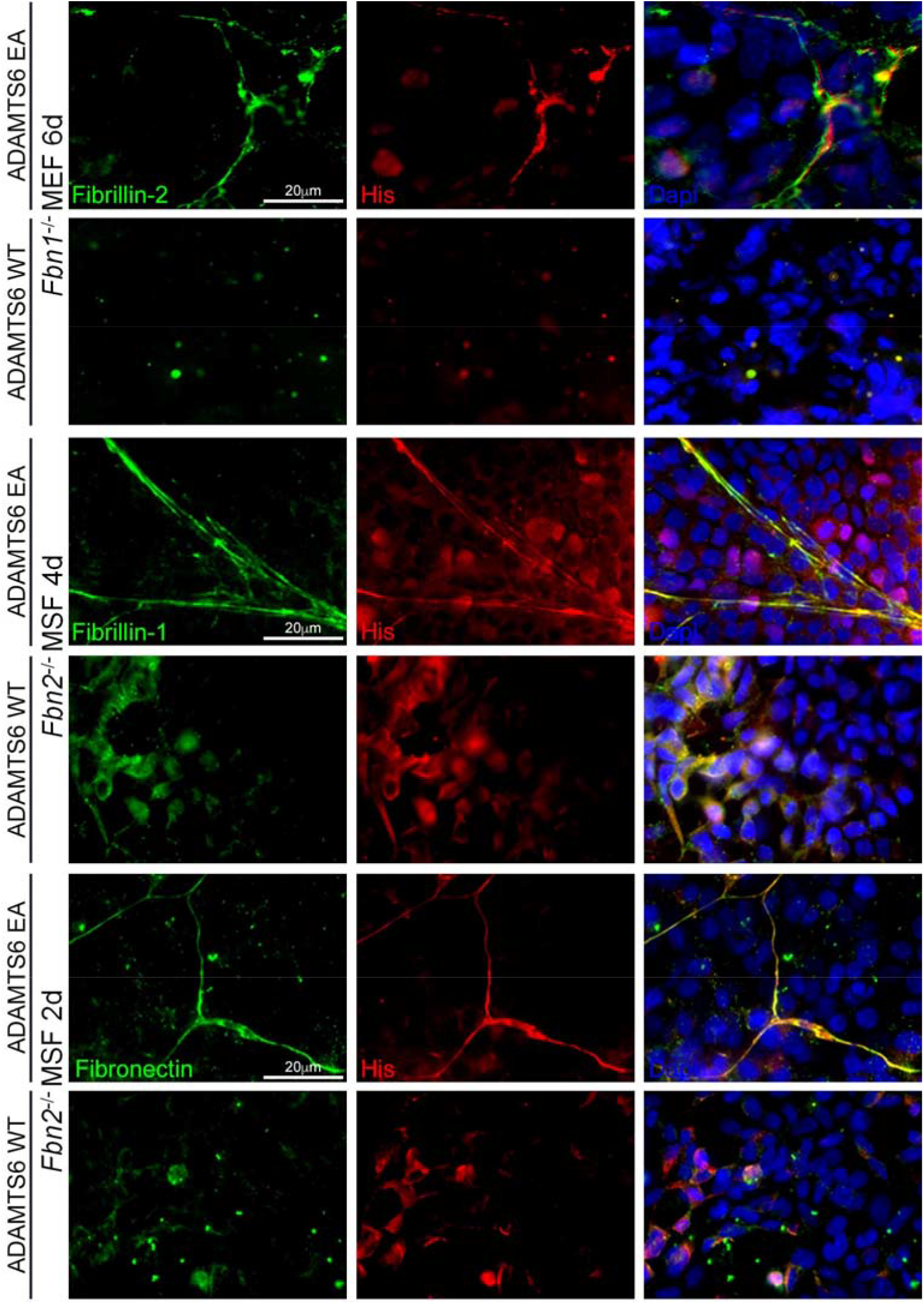
Absence of fibrillin-2, fibrillin-1 and fibronectin microfibrils in the presence of active ADAMTS6. *Fbn1*^-/-^ mouse embryo fibroblasts (MEF), which make fibrillin-2 but not fibrillin-1, were co-cultured with human embryonic kidney (HEK) cells overexpressing ADAMTS6 or ADAMTS6 EA (inactive ADAMTS6). Fibrillin-2 (green) co-localized with histidine (His)-tagged ADAMTS6 EA (red), but no microfibrils were seen in the presence of ADAMTS6 (red). Similarly, *Fbn2*^-/-^ mouse skin fibroblasts (MSF), which make fibrillin-1 but not fibrillin-2, were co-cultured with HEK cells overexpressing ADAMTS6 or ADAMTS6 EA. Fibrillin-1 microfibrils (green) co-localized with His-tagged ADAMTS6 EA (red), but fibrillin-1 microfibrils were absent in the presence of active ADAMTS6 (red) after 4 days culture. Fibronectin also co-localized with ADAMTS6 EA, but fibronectin fibrils were absent in the presence of active ADAMTS6. Nuclei are stained with DAPI

Since fibrillin microfibrils contain additional components beside fibrillins (45–48), which could be responsible for their binding to ADAMTS6, we next asked whether purified ADAMTS6 constructs bound directly to purified fibrillin-2 in binary interaction assays. ADAMTS proteases typically bind to their substrates/ECM via interactions of their C-terminal ancillary domains (49). Biacore analysis showed that C-terminal ADAMTS6 constructs (ADAMTS6-Ct, ADAMTS6-S4TSR and ADAMTS6-4TSR) bound the C-terminal half of fibrillin-2 (fibrillin-2-Ct) (Figure 5A-C, Table 1). Since all 4TSR-array-containing fragments bound to fibrillin-2-Ct, but ADAMTS6-TCS did not (data not shown), we conclude that the binding region of ADAMTS6 was located in the C-terminal TSR array. In reciprocal Biacore analysis using ADAMTS6-Ct as the immobilized ligand, binding to both the N- and C-terminal halves of fibrillin-2 was observed (Figure 5D, Table 2). Comparable Kd values of 43 nM for fibrillin-2-Nt and 80 nM for fibrillin-2-Ct suggested that the ADAMTS6 binding site on fibrillin-2 may lie in the overlapping region of the two fragments between cbEGF22 and cbEGF24, or alternatively, two or more binding sites in each half with similar affinities. Consistent with prior work showing full-length ADAMTS6 binding to fibrillin-1 (27), ADAMTS6-Ct bound to fibrillin-1 N- and C-terminal halves (Figure 5 – figure supplement 1A-B), although binding was much stronger to the N-terminal half (Figure 5 – figure supplement 1C). In summary, these binary interactions demonstrated that ADAMTS6 bound directly to both fibrillin-1 and fibrillin-2.

**Figure 5:**
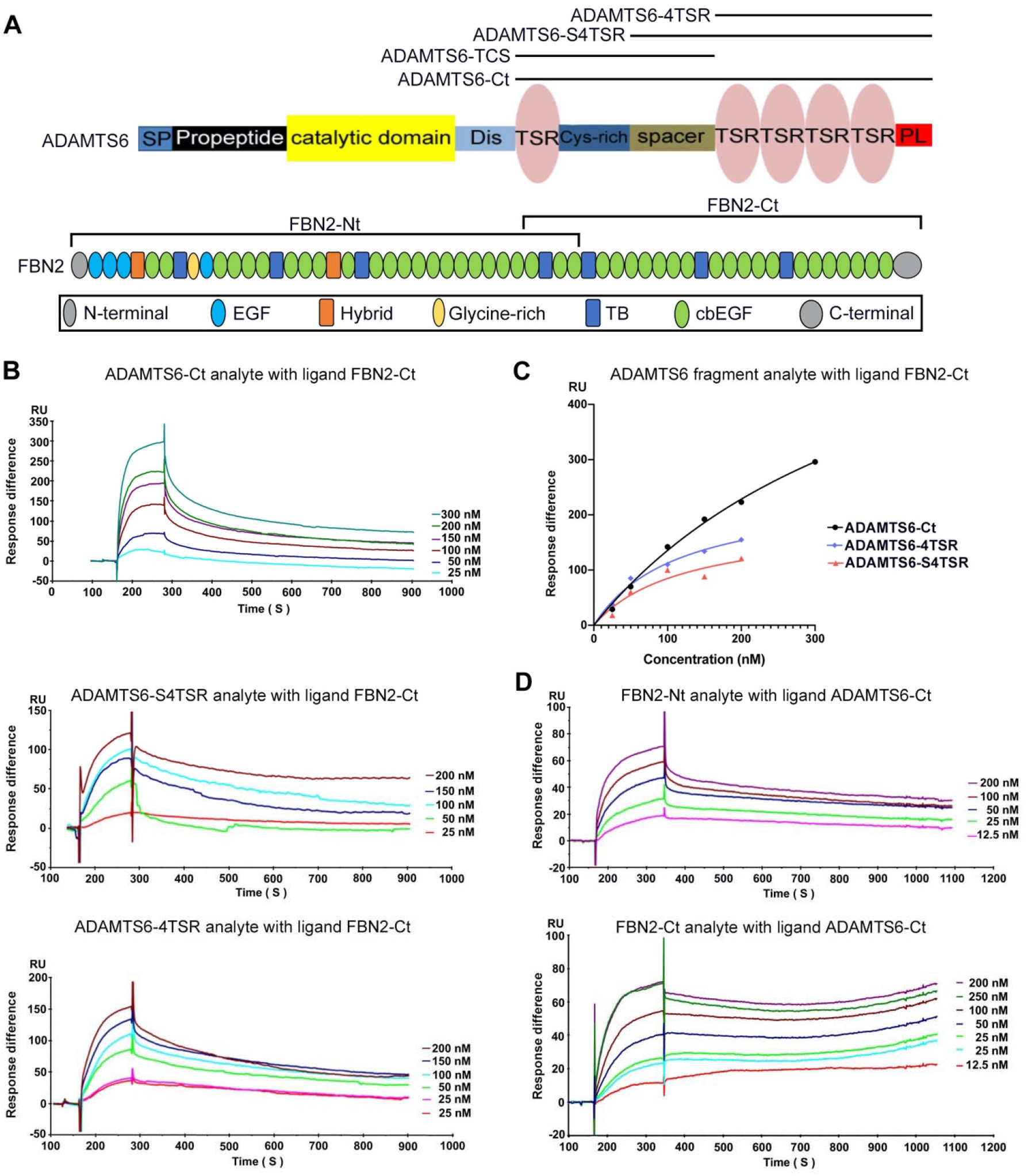
ADAMTS6 binds directly to fibrillin-2. **(A)** Domain structures of ADAMTS6 and fibrillin-2 and the recombinant protein constructs used in the present work. **(B-C**) Biacore analysis shows dose-dependent binding curves for the ADAMTS6 C-terminal constructs ADAMTS6-Ct, ADAMTS6-4TSR and ADAMTS6-S4TSR against immobilized FBN2-Ct **(B)**, and comparative binding characteristics of the constructs **(C)**. **(D)** A reciprocal Biacore analysis using immobilized ADAMTS6-Ct shows that fibrillin-2-Nt and fibrillin-2-Ct used as the analyte each bound strongly to ADAMTS6-Ct.

**Table 1.**
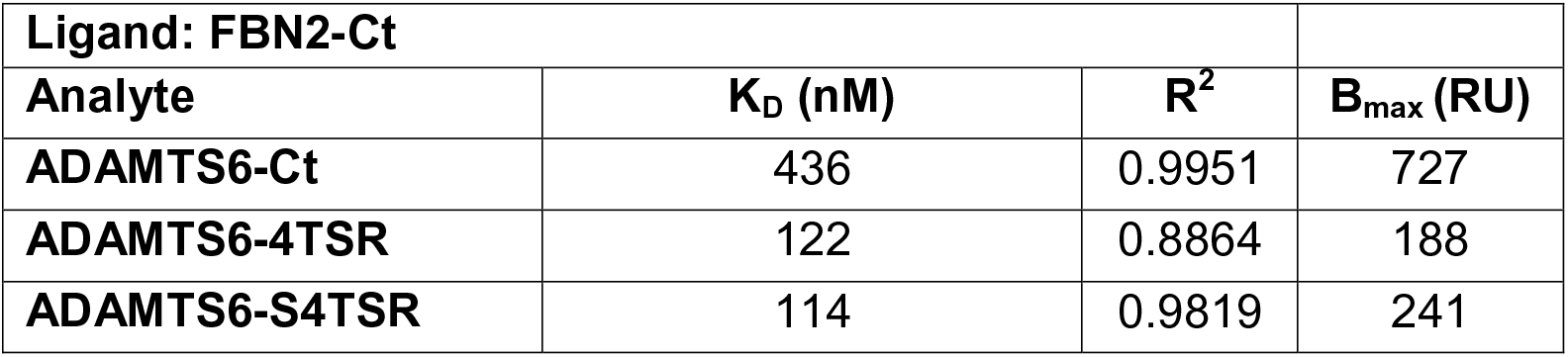
Kinetic data for ADAMTS6 construct binding to FBN2-Ct.

**Table 2.** Kinetic data for FBN2 construct binding to ADAMTS6-Ct.

| **Ligand: ADAMTS6-Ct** |  |  |  |
| --- | --- | --- | --- |
| **Analyte** | **K<sub>D</sub> (nM)** | **R<sup>2</sup>** | **B<sub>max</sub> (RU)** |
| **FBN2-Nt** | 43 | 0.9972 | 85 |
| **FBN2-Ct** | 80 | 0.9933 | 100 |

**Table 3.** Putative sites of fibronectin cleavage by ADAMTS6.

| Annotated Sequence | Number of PSMs | Wild type abundance | EA abundance |
| --- | --- | --- | --- |
| [A<sup>292</sup>].V<sup>293</sup>YQPQHPQPPPYGHCVTDSGVVYSVGMQWLK.[T] | 2 | 1.38E+06 | 0 |
| [Y<sup>294</sup>].Q<sup>295</sup>PQPHQPPPYGHCVTDSGVVYSVGMQWLK.[T] | 1 | 6.99E+05 | 0 |
| [P<sup>462</sup>].M<sup>463</sup>AAHEEICTTNEGVMYR.[I] | 2 | 1.22E+06 | 0 |
| [A<sup>464</sup>].A<sup>465</sup>HEEICTTNEGVMYR.[I] | 1 | 4.71E+05 | 0 |
| [G<sup>567</sup>].T<sup>568</sup>FYQIGDSWEK.[Y] | 1 | 1.18E+06 | 0 |
| [F<sup>750</sup>].R<sup>751</sup>VEYELSEEGDEPQYLDLPSTATSVNIPDLLPGRK.[Y] | 1 | 3.33E+05 | 0 |
| [K].YIVNVYQISEDGEQS<sup>800</sup>. [L<sup>801</sup>] | 1 | 8.57E+05 | 0 |
| [Y<sup>1248</sup>].T<sup>1249</sup>VKDDKESVPISDTIIPAVPPPTDLR.[F] | 1 | 1.21E+06 | 0 |
| [Y<sup>1613</sup>].A<sup>1614</sup>QNPSGESQPLVQTAVTTIPAPTDLK.[F]<sup>A</sup> | 1 | 6.32E+05 | 0 |
| [P<sup>2021</sup>].F<sup>2022</sup>VTHPGYDTGNGIQLPGTSGQQPSVGQQMIFEEHGFRR.[T] | 2 | 1.58E+06 | 0 |
| [K].VREEVVTVGNSVNEG<sup>2197</sup>. [L<sup>2198</sup>]<sup>B</sup> | 1 | 1.56E+06 | 0 |
| [G<sup>2197</sup>].L<sup>2198</sup>NQPTDDSCFDPTYTVSHYAVGDEWER.[M]<sup>B</sup> | 1 | 6.82E+06 | 0 |

PSMs, peptide spectrum matches.
<sup>A</sup>Peptide that is unique to the FN1-8 isoform.
<sup>B</sup>Peptides whose termini identify the same cleavage site, G<sup>2197</sup>-L<sup>2198</sup>.

### N-terminomics identification and orthogonal validation of an ADAMTS6 cleavage site in fibrillin-2

We investigated whether increased fibrillin-2 staining in *Adamts6*-deficient hindlimbs and lack of fibrillin-2 microfibrils *in vitro* in the presence of ADAMTS6 indicated a role in fibril proteolysis or assembly. Direct binding of ADAMTS6 constructs to fibrillin-2 protein, ADAMTS6 EA co-localization with fibrillin-2 microfibrils, loss of fibrillin-2 microfibrils in vitro in the presence of wild type ADAMTS6, and increased fibrillin-2 staining in *Adamts6*-deficient hindlimbs suggested that fibrillin-2 was an ADAMTS6 substrate. Furin site-optimized ADAMTS10 was previously shown to cleave fibrillin-2, and *Adamts10*^-/-^ mice had accumulation of fibrillin-2 microfibrils in the eye and skeletal muscle (29, 30). Therefore, HEK293F cells stably expressing the N- or C-terminal halves of fibrillin-2 were transfected with either ADAMTS6 or ADAMTS6 EA and the serum-free conditioned medium was collected for Terminal Amine Isotopic Labeling of Substrates (TAILS), an N-terminomics approach for identifying protease substrates and cleavage sites (50, 51), which was previously applied for identification of substrates of other ADAMTS proteases (52) (Figure 6A). Proteins were labeled with stable isotopes of formaldehyde (natural (CH_2_O)/light isotope applied to the ADAMTS6-containing medium or isotopically heavy (^13^CD_2_O), applied to the ADAMTS6 EA-containing medium). The ensuing reductive dimethylation labels and blocks free protein N-termini as well as lysine sidechains. Labeled proteins from each pair of ADAMTS6/ADAMTS6 EA digests were combined, further digested with trypsin to obtain peptides and mixed with hyperbranched polyglycerol aldehyde polymer which binds to free N-termini of tryptic peptides (Figure 6A), enriching peptides having blocked and labeled N-termini for liquid chromatography tandem mass spectrometry (LC-MS/MS). Following LC-MS/MS, a targeted search for differentially abundant fibrillin-2 peptides with isotopically labeled N-termini having a statistically significant light:heavy ratio revealed a putative cleavage site at the Gly^2158^-His^2159^ peptide bond in a linker between TGFβ binding-like domain 6 and epidermal growth factor (EGF) repeat 32 in the C-terminal half of fibrillin-2 (Figure 6B-F). The sequence of the indicator peptide and presence of an N-terminal label was confirmed with high confidence by the MS2 spectrum (Figure 6C). Quantification of the isotoptically-labeled peptides’ ion abundance in samples showed a considerable excess of this peptide in the presence of active ADAMTS6 (Figure 6D-E). Western blot of the medium from these experiments for orthogonal validation of the cleavage showed that ADAMTS6 cleaved the C-terminal half of fibrillin-2 (Figure 6G), but not the N-terminal half (data not shown). The cleavage products of 100 kDa and 75 kDa matched the predicted cleavage fragments in size and their sum matched the expected mass of the FBN2-Ct construct (175 kDa). These findings strongly suggested that fibrillin-2 is an ADAMTS6 substrate which could be relevant to the profound skeletal defects observed in *Adamts6*^-/-^ mice. Importantly, the Gly^2158^-His^2159^ cleavage site is located between two folded, disulfide-bonded domains, predicting separation of the resulting fragments (as opposed to cleavage sites within fibrillin-2 domains that would remain linked by disulfide bonds). The cleavage can potentially interfere with microfibril assembly, which is reliant on multimerization via C-terminal interactions (i.e., downstream of the cleavage site) (53).

**Figure 6:**
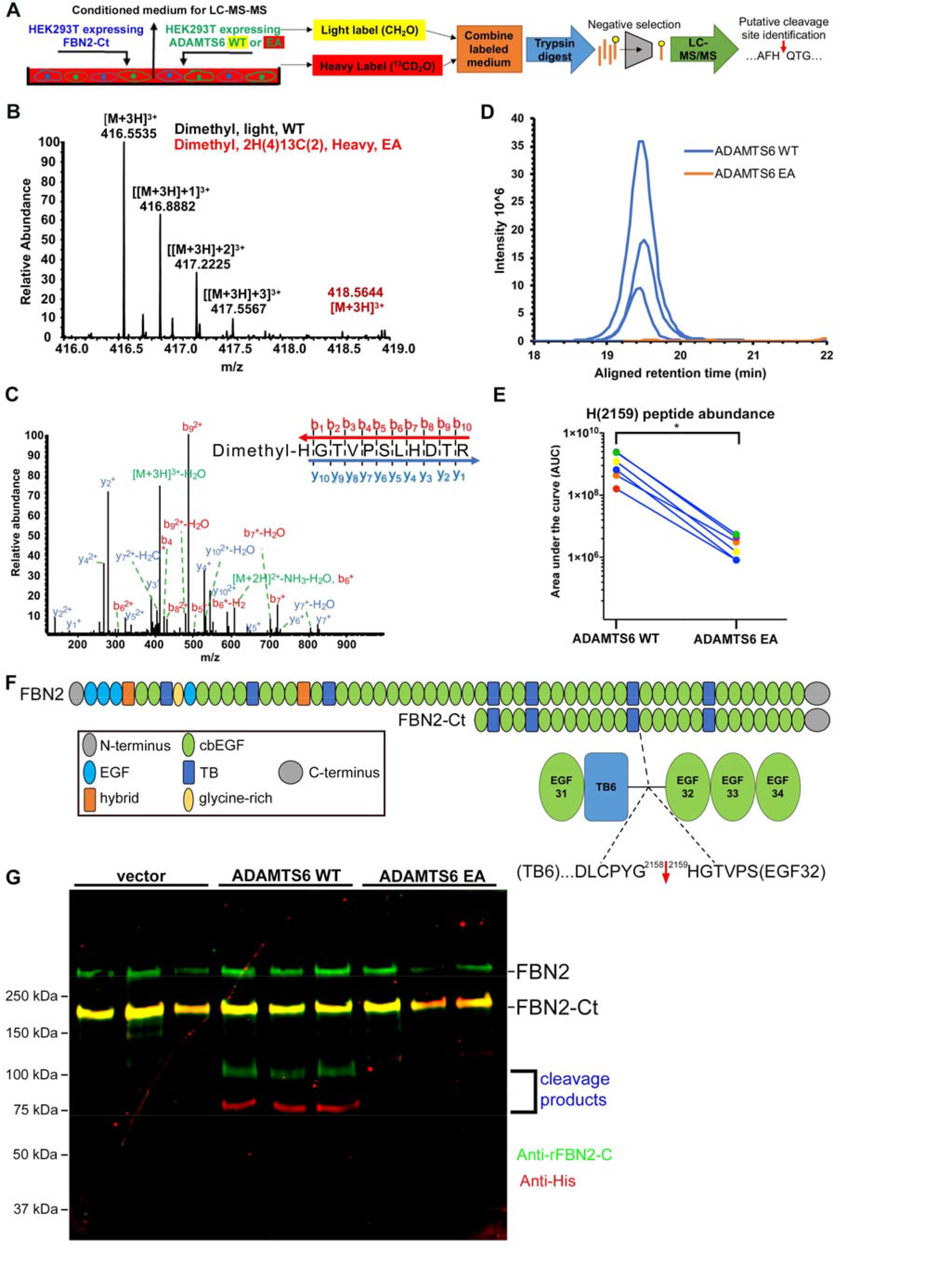
Fibrillin-2 cleavage by ADAMTS6 and identification of the cleavage site using N-terminomics. **(A)** Schematic of the experimental approach. Proteins from conditioned medium of co-cultures of HEK293F cells stably expressing FBN2-Ct and cells expressing either ADAMTS6 WT or ADAMTS6 EA (inactive) were labeled by reductive dimethylation using stable formaldehyde isotopes and analyzed by LC-MS/MS in the TAILS workflow described in detail in the Methods section. **(B)** MS1 chromatogram of the light and heavy isotope-labeled parent ions representing fibrillin-2 peptides from ADAMTS6 WT (black) and ADAMTS6 EA-containing medium (red) that were used for quantitation. (**C**) Annotated MS2 spectrum of the light (ADAMTS6-generated) dimethyl peptide, showing b-(N-terminus preserved) and y-type (C-terminus preserved) ions generated by amide bond cleavage during collisional-induced dissociation that were used to derive the fibrillin-2 peptide sequence indicated at top right. **(D)** Retention time-aligned extracted ion chromatographs (EICs) comparing abundance of the light dimethyl-labeled HGTVPSLHDTR peptide (blue) in ADAMTS6 WT medium and isotopically heavy dimethyl-labeled peptide (orange) in ADAMTS6 EA medium from the 3 TAILS replicate experiments. **(E)** The area under the EICs was quantified and comparison of ion abundance is shown in a dumbbell plot (from 3 TAILS and 3 pre-TAILS samples). Statistical significance was determined using a two-tailed, paired Student t-test, * indicates *P*-value < .05. (**F)** Domain structure of fibrillin-2 and the C-terminal construct FBN2-Ct showing the location of the cleaved peptide bond Gly^2158^-His^2159^ in the linker between TB6 and EGF32. **(G)** Orthogonal validation of fibrillin-2 cleavage by ADAMTS6 using western blot analysis of the conditioned medium from A, shows distinct molecular species (100 kDa and 75 kDa) reactive with anti-fibrillin-2-Ct antibody (green, N-terminal fragment of fibrillin-2-Ct) and C-terminal anti-His_6_ antibody (red, C-terminal fragment of fibrillin-2-Ct), respectively, obtained in the presence of ADAMTS6, but not ADAMTS6 EA, indicative of fibrillin-2-Ct cleavage. The green band of ∼350 kDa is endogenous fibrillin-2 produced by HEK293T cells. The yellow band at ∼175 kDa indicates overlapping anti-His_6_ and anti-fibrillin-2 Ct antibody staining of fibrillin-2-Ct. Cells transfected with empty vector were used to obtain control medium.

### ADAMTS6 cleaves fibrillin-1 and fibronectin

Because ADAMTS6 also bound directly to fibrillin-1, TAILS was applied in a similar approach as above to determine if ADAMTS6 cleaved fibrillin-1 (Figure 6 – figure supplement 1A). A targeted search for fibrillin-1 peptides with labeled N-termini revealed a putative cleavage site at the R^516^-A^517^ peptide bond in a linker between epidermal growth factor (EGF) repeats 6 and 7 in the N-terminal half (Figure 6 – figure supplement 1B). The peptide sequence and presence of an N-terminal label was confirmed with high confidence by the MS2 spectrum and quantification of the isotopically labeled peptides’ ion abundance showed a considerable excess of this peptide in the presence of ADAMTS6 (Figure 6 – figure supplement 1C-E).

To determine whether ADAMTS6 cleaved fibronectin, ADAMTS6 or ADAMTS6 EA-expressing stable cell lines were plated on human fibronectin-coated plates and the serum-free medium from these cultures was collected 24 h later. By comparing the ratio of light:heavy dimethyl-labeled peptides, TAILS revealed several potential cleavage sites in the medium from the ADAMTS6 sample (Figure 6 – figure supplement 2A). As an example, annotated spectra of two of the peptides that identify the same cleaved peptide bond, and their extracted ion chromatograms illustrate the lack of the corresponding peaks in medium of the ADAMTS6 EA samples, indicating fibronectin cleavage solely in the presence of catalytically active ADAMTS6 (Figure 6 – figure supplement 2B-E).

### Fibrillin-2 reduction, but not Fibrillin-1 reduction reverses morphogenetic and molecular defects in *Adamts6^-/-^* mice

To determine whether reduced fibrillin-2 and/or fibrillin-1 cleavage had a significant role in the observed anomalies, fibrillin-2 was reduced genetically in *Adamts6*^-/-^ mice. *Fbn2* haploinsufficiency dramatically restored the external craniofacial and hindlimb morphology of *Adamts6* mutants as well as the maturity, length and shape of skeletal components of the hindlimb and axial skeleton (Figure 7A,B). Specifically, alizarin red and alcian blue-stained skeletal preparations demonstrated reversal of bone shortening and improved ossification of hindlimb long bones and ribs, and an appropriately segmented sternum with a normal xiphoid process (Figure 7A, B, Figure 7 – figure supplement 1, Figure 7 – figure supplement 2A). In addition, cleft secondary palate which occurred with a high incidence in *Adamts6*^-/-^ embryos, was not observed in *Adamts6*^-/-^;*Fbn*2^+/-^ embryos (Figure 7 – figure supplement 2B). *Adamts6*^-/-^; *Fbn*2^+/-^ growth plate histology was comparable to wild type and fibrillin-2 staining in the hindlimb showed reduced intensity comparable to *Adamts6*^-/-^ hindlimb (Figure 7C, Figure 7 – figure supplement 3). Aggrecan, cartilage link protein and Sox9 staining intensity in *Adamts6*^-/-^;*Fbn*2^+/-^ femur growth plates were restored to comparable levels as in wild type limbs (Figure 7D). In contrast *Adamts6*^-/-^ mice with ∼80% reduction in fibrillin-1 levels (i.e., *Adamts6*^-/-^;*Fbn1*^mgR^/^mgR^ embryos) (Figure 7 – figure supplement 4) had no amelioration of limb defects, suggesting a selective impact of ADAMTS6 on fibrillin-2 abundance in limb microfibril proteostasis.

**Figure 7:**
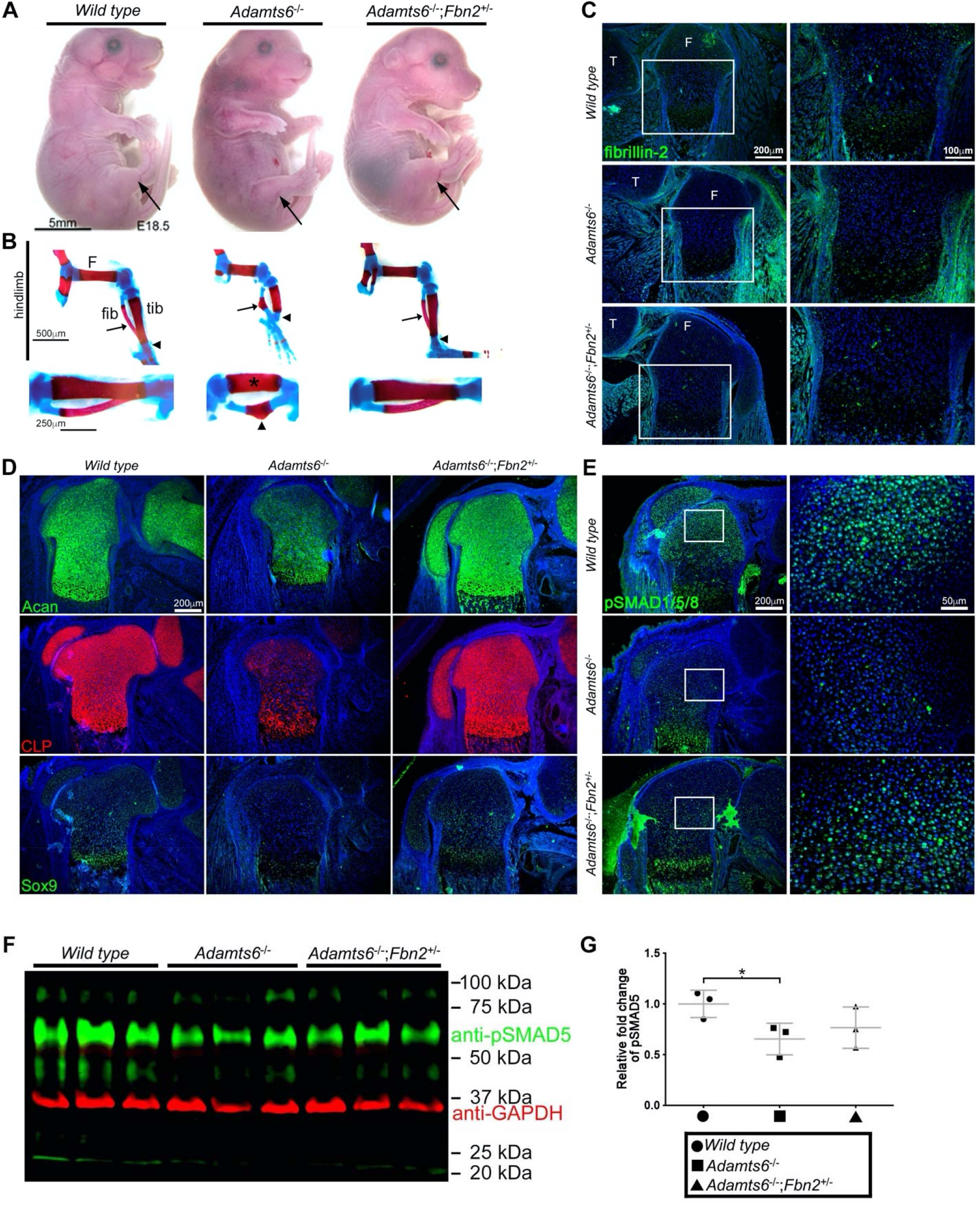
Genetic reversal of *Adamts6* mutant skeletal anomalies by *Fbn2* haploinsufficiency. **(A-B)** Deletion of one *Fbn2* allele reverses limb dysmorphology in E18.5 *Adamts6*^-/-^ embryos, specifically, externally evident limb segment dimensions and reversal of rotational anomaly (arrow) (**A**), normal ossific centers and overall hindlimb skeletal structure, reversal of internal rotation (arrow) and bending of tibia (upper arrowheads) and restores normal tibial (asterisk) and fibular (lower arrowhead) length and alignment. **(C)** Fibrillin-2 staining is reduced to wild type levels in *Adamts6*^-/-^;*Fbn2*^+/-^ distal femoral perichondrium. **(D)** Restitution of aggrecan, cartilage link protein (CLP) and Sox9 staining in *Adamts6*^-/-^;*Fbn2*^+/-^ distal femoral cartilage. **(E)** Reduced pSMAD1/5/8 staining in *Adamts6*^-/-^ femur, as compared to wild type control, is restored in *Adamts6*^-/-^;*Fbn2*^+/-^ femur. **(F)** Western blot analysis shows reduced pSMAD5 (green; 58 kDa) in *Adamts6*-deficient E18.5 hindlimb lysates. Anti-GAPDH (red, 37 kDa) was used as a loading control. **(G)** pSMAD5 in (F) was quantified after normalization to GAPDH loading control. n= 3. *p ≤0.05.

Consistent with the previously defined role of fibrillin-2 in regulating BMP signaling, *Adamts6*^-/-^ cartilage had reduced pSmad5 staining compared to wild type, which was restored to wild type levels in *Adamts6*^-/-^;*Fbn*2^+/-^ hindlimbs (Figure 7E-G, Figure 7 – figure supplement 5). In contrast, no change in TGFβ signaling, as measured by pSMAD2 immunostaining and western blot, occurred in *Adamts6*^-/-^ cartilage (Figure 7 – figure supplement 6).

## Discussion

Regulated ECM proteostasis is required for ensuring proper adult tissue function (mitigating the risk of organ fibrosis, for example), but presumably also during transition from the early to late embryo ECM, and may be a major determinant of adaptation to the dramatically different postnatal mechanical and regulatory landscapes. Because of the immense complexity of molecules and networks that comprise the ECM, as well as numerous secreted proteases, we hypothesized that selective proteolysis arising from co-evolution of specific proteases with preferred ECM substrates, may have a crucial role in ECM proteostasis in specific contexts of embryonic development. ECM proteostatic phenomena that are now well described include fibrillar collagen breakdown by MMPs and versican turnover by ADAMTS proteases (34, 54–56), but macromolecular complexes formed by homologous proteins with varying stoichiometry in the embryo and adult present an interesting challenge in regard to selectivity and mechanisms.

Here, using a combination of genetic, biochemical and in vitro approaches, we demonstrate the role of ADAMTS6 and ADAMTS10 in fine-tuning of fibrillin microfibrils. Although prior work linked ADAMTS10 to genetic conditions that also resulted from *FBN1* mutations, the existence of a closely related homolog, ADAMTS6, raised questions about the role of transcriptional adaptation in this protease pair, their overlapping functions on a organ, tissue and molecular scale, and the identity of their specific targets. Resolution of these questions provides a conceptual precedent for further elucidation of embryonic ECM proteostasis, about which little is known relative to cellular homeostatic mechanisms in the embryo.

In the context of mechanisms that would ensure fibrillin-1 predominance postnatally, prior RNA in situ hybridization analysis had demonstrated dramatic reduction of *Fbn2* mRNA expression postnatally, with *Fbn1* expression continuing (13) and corresponding to this, little fibrillin-2 is detected immunohistochemically or by proteomics in adult mouse tissues (46, 53, 57–60). Thus, differential transcription of fibrillin genes favors reduced fibrillin-2 synthesis postnatally, and together with dominance of *Fbn1* expression, was thought to underlie the prevalence of fibrillin-1 microfibrils in juvenile and adult mice. Fibrillin-2 was present postnatally in only a few locations, and that some tissue microfibril bundles had a core of fibrillin-2 microfibrils surrounded by abundant fibrillin-1, suggested that fetal microfibrils formed an inner core to which postnatal microfibrils (comprising fibrillin-1) were added (46, 61). Previous work also suggested that fibrillin-2 epitope availability was masked postnatally by such overlay of fibrillin-1. Specifically, it was shown that fibrillin-2 staining in postnatal tissues such as perichondrium was enhanced by digestion with collagenase, that microfibrils treated with chaotropic agent had enhanced reactivity to fibrillin-2 antibodies, and that *Fbn1* null juvenile mice, which die by 2 weeks of age, had robust fibrillin-2 staining (61). Thus, it was thought that the fibrillin isoform content of microfibrils is substantially, if not solely, determined by the level of transcription of the respective genes. The fate of fibrillin-2 microfibrils produced during the embryonic period, the possible existence of specific proteolytic mechanisms to reduce their abundance, and a deleterious impact of fibrillin-2 overabundance had not been previously considered and no animal models with excess fibrillin-2 were available.

The present work demonstrates that two homologous fibrillin-2 degrading proteases work collaboratively, but in distinct ways, to support prevalence of fibrillin-1 in microfibrils postnatally. For example, ADAMTS10 is innately resistant to activation by furin, with only a small proportion converted to an active protease. ADAMTS10 promoted fibrillin-1 microfibril assembly (28), consistent with the observation that recessive *ADAMTS10* mutations and dominantly inherited *FBN1* mutations each led to Weill-Marchesani syndrome (23, 62). ADAMTS10 was therefore thought to function akin to several ADAMTS-like proteins, which lack catalytic activity, and accelerate biogenesis of fibrillin-1 containing microfibrils (25, 63–68). ADAMTS10, if furin-activated by introduction of a furin cleavage site, was recently demonstrated to be proteolytically active against fibrillin-2 (30), which raised the possibility that its nearest homolog, ADAMTS6 may cleave fibrillin-2. The present work shows that ADAMTS6, whose zymogen is efficiently activated by furin (27), cleaves both monomeric fibrillin-1 and fibrillin-2, has identified specific sites of cleavage using TAILS and validated proteolysis biochemically in the case of the fibrillin-2 site. Fibrillin microfibril proteolysis was supported by loss of fibrillin-1 and fibrillin-2 microfibrils assembled by *Fbn1*^-/-^ and *Fbn2*^-/-^ fibroblasts, respectively, in the presence of active ADAMTS6. Although we are unable to purify full-length ADAMTS6 protease, and thus, cannot exclude the possibility that ADAMTS6 acts via activation of another protease in the cell culture system that was used, the direct binding of ADAMTS6 to the fibrillins in a binary interaction assay strongly supports the likelihood that it cleaves them directly. The significance of the loss of fibrillin-2 proteolysis and consequent accumulation resulting from *Adamts6* inactivation was demonstrated by dramatic reversal of the observed skeletal and palate defects after *Fbn2* haploinsufficiency in the *Adamts6* mutant. The specificity for fibrillin-2 was further established by lack of a comparable effect after reduction of fibrillin-1. While we cannot exclude the likelihood of accumulation of additional ADAMTS6 proteolytic targets in the skeleton, such as LTBP-1, which it was previously shown to cleave (27), it is clear that fibrillin-2 proteolysis by ADAMTS6 is necessary for proper skeletogenesis.

The present work clarifies the impact of transcriptional adaptation and functional cooperativity, which have not been hitherto analyzed in detail in ECM and ECM proteases. In contrast to *Adamts6*-deficient mice, *Adamts10*^-/-^ mice have subtle skeletal defects. However, persistent fibrillin-2 fibrils were noted in *Adamts10*-deficient eyes, specifically in the zonule and vitreous, as well as in skin and muscle (29, 30). Although furin-activated ADAMTS10 cleaved fibrillin-2, we concluded that ADAMTS10 may have a minor role in proteolytic degradation of fetal fibrillin-2 microfibrils in the eye and elsewhere (30). The observed upregulation of *Adamts6* mRNA in *Adamts10*-deficient embryos shows it could compensate for *Adamts10* by transcriptional adaptation (38, 39) and may also explain the mild phenotype of *Adamts10* targeted mice (30) including the modest accumulation of fibrillin-2 we observed here. However, further reduction of bone length after introducing an *Adamts10* mutant allele into *Adamts6*^-/-^ mice, where transcriptional adaptation does not occur, suggests that ADAMTS6 and ADAMTS10 cooperate in skeletal development. The phenomenon of cooperative fibrillin-2 proteolysis is supported by the activity of both proteases against this substrate. While *Adamts*6 does compensate for *Adamts10*-deficiency, ADAMTS6 and ADAMTS10 do not proteolytically activate/modify each other suggesting that they work cooperatively in parallel rather than in series, using the analogy of electrical circuits. Prior work showed that transcriptional adaptation is also operational in genes encoding paired homologous proteases ADAMTS7 and ADAMTS12 (69), and indeed, may be a widespread phenomenon in this protease family, which has arisen largely from gene duplication (2).

In addition to mechanical effects of excess perichondrial fibrillin-2 on cartilage that could affect skeletal development, reduced fibrillin-2 proteolysis could lead to dysregulated sequestration and release of growth factors of the TGFβ superfamily, since fibrillin-2 can bind to numerous superfamily members including several BMPs and GDFs (18, 70, 71). In this regard, prior work has shown that fibrillin microfibrils do not have a role in activation of the BMPs, but primarily affect local concentrations and release, which may be mediated in part via BMP proteolysis by MMPs (72, 73). In contrast, specific fibrillin proteolytic events have not been implicated in control of local BMP/GDF activity. In the present analysis, the microfibril changes in the perichondrium were associated with changes in the underlying cartilage, in further support of phenotype modulation across the perichondrium-cartilage boundary (74–76). Furthermore, prior work has shown that combinations of TGFβ superfamily members strongly promote maintenance of expression of typical cartilage-specific genes (77, 78). This previous work, together with decreased pSmad5 levels (indicative of reduced BMP signaling) may explain reduction of Sox9, aggrecan and link protein that we observed. The resulting reduced cartilage proteoglycan may also result in softer cartilage that could be susceptible to bending moments as limb muscles develop, potentially explaining the angulated chondroepiphyses.

Accumulation of fibrillin-2 microfibrils resulting from combined ADAMTS6 and ADAMTS10 deficiency may excessively sequester multiple BMPs and GDFs in the perichondrium, leading to impaired BMP signaling in the underlying cartilage. Consistent with this possibility, *Fbn2*-deficent mice have increased BMP signaling (18). Fibrillins interact with the related latent TGFβ-binding proteins and MAGP1, a fibrillin-binding partner shown to accumulate in the *Adamts6* mutant mice also regulates the cellular microenvironment via growth factor binding (79, 80). Moreover, ADAMTS6-mediated proteolysis of cell-surface heparan-sulfate proteoglycans (27) could also influence the skeletal microenvironment. Altogether, complex effects on a variety of soluble morphogens and growth factors may ensue downstream of the observed ECM changes, but the genetic evidence from our studies unequivocally indicates that fibrillin-2 accumulation is a central mechanism underlying the observed skeletal defects. These findings highlight the importance of specific proteolytic mechanisms in ECM proteostasis and their importance for cellular regulation.

## Materials and Methods

### Transgenic mice

*Adamts6*^b2b2029Clo^ (RRID:MGI:5487287), *Adamts10*^tm1Dgen^ (MGI:6355992), *Fbn1*^mgR^ (*Fbn1*^tm2Rmz^; MGI:1934906) and *Fbn2*^tm1Rmz^ (RRID:MGI:3652417) mice were previously described (17, 30, 31, 81, 82) and maintained on a C57BL/6 background. All mouse experiments were approved by the Cleveland Clinic Institutional Animal Care and Use Committee (protocol 2018-2450).

### Histology and immunofluorescence microscopy

Mouse forelimbs and hindlimbs were fixed with 4% paraformaldehyde (PFA) in PBS at 4 °C for 48 hrs. 7μm sections were used for histochemistry (alcian blue or RGB trichrome stain (83)) or for indirect immunofluorescence. The primary antibodies (Supplemental Table 1) were followed by secondary goat anti-mouse or goat anti-rabbit antibody (A11004 or A11008; 1:400; Invitrogen, Thousand Oaks, CA) treatment. Prior to immunofluorescence, citrate antigen retrieval, i.e., immersion of slides in citrate-EDTA buffer (10 mM/l citric acid, 2 mM/l EDTA, 0.05% v/v Tween-20, pH 6.2) and microwaving for 4 intervals of 1.5 min at 50% power in a microwave oven with 30 s intervals between heating cycles was utilized. In addition, hyaluronidase treatment of sections (H-2251; 0.2% in PBS; Sigma) was used prior to fibrillin-2 and fibrillin-1 immunostaining. After deparrifinization and rehydration, sections were stained with 1% alican blue 8GX (A3157; Sigma) in 3% acetic acid (pH 2.5) for 10 min, rinsed in tap water, counter-stained in nuclear fast red (J61010; Alfa Aesar) for 1 min, rinsed in tap water prior to dehydration and mounting as previously described (84). Alcian blue-stained sections used for hypertrophic chondrocyte zone length were masked during data collection and measured along the midline of the femoral and tibial growth plate using NIH Fiji software (85) as previously reported (35). For each embryo, data was generated from 3 separate sections, spaced at least 25 μm apart and averaged. RGB-trichrome staining was modified as previously described (83). Briefly, after de-parrafinization and rehydration, the sections were stained with 1% alcian blue 8GX (A3157; Sigma) in 3% acetic acid (pH 2.5) for 20 min, rinsed in tap water, 1% fast green FCF (F-99; Fisher Scientific) for 20 min, rinsed in tap water, 0.1% sirius red in saturated picric acid (26357-02; EMS) for 60 min, rinsed in 2 changes of 1% acetic acid prior to dehydration in 100% ethanol and xylene. TUNEL assay (11684795910; Sigma) was performed as described previously (86). Images were obtained using an Olympus BX51 microscope with Leica DFC 7000T camera and Leica Application Suite V4.6 software (Leica, Wetzlar, Germany). Histological sections were masked during data collection and quantified utilizing NIH Fiji software (85).

Alizarin red-alcian blue stained skeleton preparations were performed as described (87). Briefly, skinned, eviscerated mice were fixed in 80% ethanol for 24 h, dehydrated in 95% ethanol for 24 h and acetone for 48 h, stained (0.1% alizarin red S, 0.3% alcian blue, 1% glacial acetic acid in 95% ethanol) for 48 h, cleared in 95% ethanol for 1 h, muscle tissue was gently removed with forceps, and the preparations were cleared in a series of increasing (20 to 80%) glycerol/1% KOH ratio until storage in 100% glycerin and photography (Leica MZ6; Insight Spot software camera and software). Crown to rump or individual bone length measurements were masked during data collection utilizing NIH Fiji software.

### RNA in situ hybridization (ISH) and β-Gal staining

Hindlimbs from *Adamts10*^-/-^ embryos were fixed and β-galactosidase-stained as previously described (88), followed by paraffin embedding and 10 μm sections were obtained. *Adamts6* ISH was performed using RNAScope (Advanced Cell Diagnostics, Newark, CA) as described (89). Briefly, 6 μm sections were deparaffinized and hybridized to a mouse *Adamts6* probe set (428301; Advanced Cell Diagnostics) using a HybEZ^TM^ oven (Advanced Cell Diagnostics) and the RNAScope 2.5 HD Detection Reagent Kit (322360; Advanced Cell Diagnostics) and counterstained with eosin.

### Cell culture

HEK293T cells were purchased from ATCC and maintained in Dulbecco’s Modified Eagles Medium (DMEM) supplemented with 10% fetal bovine serum (FBS), 100 U/ml penicillin and 100 μg/ml streptomycin at 37 °C in a 5% CO_2_ humidified chamber. The cells were transiently transfected with ADAMTS6 WT or ADAMTS6 EA expression plasmids using PEI MAX (24765; Polysciences) and were co-cultured with *Fbn1*^-/-^ mouse embryo fibroblasts (MEFs) or *Fbn2*^-/-^ mouse skin fibroblasts (MSFs) in a 1:1 ratio on 8-well culture slides (354118; Falcon). Similarily, *Adamts6*^-/-^ and wild type MEFs were plated on 8-well culture slides for immunofluorescent staining. The cells were cultured for 6 days, fixed in ice-cold 70% methanol/30% acetone for 5 min at room temperature, blocked with 5% normal goat serum in PBS for 1 h at room temperature and incubated with primary antibody (see Supplemental Table 1) overnight at 4 °C as described (90). The cells were washed 3 times with PBS for 5 min at room temperature and incubated with Alexa-Fluor labeled secondary antibodies (goat anti-mouse 568 or goat anti-rabbit 488; Invitrogen A11004, A11008, respectively, 1:400). In order to identify if fibronectin was an ADAMTS6 substrate, HEK 293T cells were transiently transfected with either ADAMTS6 or ADAMTS6 EA plasmids and seeded onto wells coated with human fibronectin as previously described (91).

### RNA isolation and Quantitative Real-Time PCR (qRT-PCR)

Mouse hindlimbs, hearts and lungs were snap-frozen and stored at −80 °C until use. Total RNA was isolated using TRIzol (15596018, Invitrogen), and 2μg of RNA was reverse transcribed into cDNA using a High-Capacity cDNA reverse transcription kit following the manufacturer’s instructions (4368814; Applied Biosystems, Foster City, CA). qRT-PCR was performed with Bullseye EvaGreen qPCR MasterMix (BEQPCR-S; MIDSCI) using an Applied Biosystems 7500 instrument. The experiments were performed with three independent biological samples and confirmed reproducibility with two to three technical replicates. *Gapdh* was used as a control for mRNA quantity. The ΔΔCt method was used to calculate relative mRNA expression levels of target genes and shown as standard error of the mean (SEM). See Supplemental Table 2 for primer sequences.

### Surface plasmon resonance analysis

The human fibrillin-2 recombinant halves (rFBN2-N and rFBN2-C, termed FBN2-Nt and FBN2-Ct in this manuscript, respectively) were purified to homogeneity (>90% purity) as described previously (21). Purified FBN2-Ct or ADAMTS6-Ct in 10 mM acetate, pH 4.0 were immobilized on a Biacore CM5 sensor chip (research grade) with the amine coupling kit following the manufacturer’s instructions (GE Healthcare). 1700 resonance units of FBN2-Ct or ADAMTS6-Ct was coupled to the chip for analysis in a Biacore 3000 instrument (GE Healthcare). The kinetics analysis was performed at 25 °C in 10 mM Hepes buffer, pH 7.4 with 0.15 M NaCl, 2 mM CaCl_2_, and 0.005% or 0.05% (v/v) surfactant P20 l/min. All the analytes were diluted in the above buffer at different μ concentrations and injected through an uncoupled control flow cell in series with the flow cell coupled with FBN2-Ct or ADAMTS6-Ct constructs. The sample injection time was 2 min for ADAMTS6 and 3 min for FBN2 analytes. The dissociation time was 10 min. 1 M ethanolamine, pH 8.5 was used for chip surface regeneration at a flow rate of 30 μl/min for 30–60 s followed by 2 min stabilization time. All data were corrected with reference to the background binding in the control flow cell. The association and disassociation curves were generated with the BIAevaluation software (version 4.0.1; GE Healthcare). The kinetic constants were calculated using the steady state affinity method.

### Site-directed mutagenesis, transient transfection, deglycosylation and western blotting

A plasmid encoding mouse ADAMTS6 with a C-terminal myc/his tag was generated previously (31) and used for site-directed mutagenesis (Q5 Site-Directed Mutagenesis Kit; E0554; New England BioLabs) to introduce Ala at Glu^404^, a classic metalloprotease inactivating mutation (ADAMTS6 EA). Plasmids were transfected into HEK293F cells using PEI MAX (24765; Polysciences) and conditioned medium was collected 48-72 h later. Aliquots of medium were analyzed by 7.5% reducing SDS-PAGE. Proteins were electroblotted to polyvinylidene fluoride membranes (IPFL00010, EMD Millipore, Billerica, MA), incubated with primary antibodies (see Supplemental Table 1) overnight at 4 °C, followed by fluorescent secondary antibody goat anti mouse or rabbit (827-08365, 926-68170; 1:5000; Li-COR Biosciences, Lincoln, NE) for 1 h at room temperature. Antibody binding was visualized using an ODYSSEY CLx infrared imaging system (LI-COR). For pSMAD5 detection, hindlimbs were placed in Ripa Buffer (ab156034; Abcam) and Complete Protease Inhibitor Cocktail (no. 4693159001; Millipore) and PhosSTOP (Millipore no. 4906845001) were added prior to homogenization (T10 basic ULTRA-TURRAX (IKA, Staufen, Germany) and ultrasonication (Qsonica, Newtown, CT, USA)( 3 × 2 s at 20% with 3 s pause). The supernantant was collected after centrifugation and 100 μg loaded on a 10% gel. Western blot band intensity was quantified utilizing NIH Fiji software (85).

### TAILS sample workflow

Serum and phenol red-free conditioned medium from cell cultures were centrifuged at 4000 rcf for 20 min at 4 °C and the supernatant was filtered through a 0.22 µM filter. The medium was concentrated 20-fold using a 3 kDa (Amicon) stirring filter. Proteins were isolated using chloroform/methanol precipitation and resuspended in 2.5 M GuHCl and 250 mM HEPES pH 7.8. Protein concentration was measured using the Bradford assay (Pierce, Thermo) to determine the volume needed for 500 µg of protein from each condition. Proteins were reduced with 10 mM dithiothreitol (DTT) for 30 min at 37 °C followed by alkylation with 20 mM N-ethylmaleimide in the dark for 20 min. The reaction was quenched by adding DTT to a final concentration of 20 mM. Proteins were labeled overnight with 40 mM light or heavy formaldehyde, which binds specifically to free N-termini and lysine residues (α and εamines, respectively) in the presence of 20 mM sodium cyanoborohydride at 37 °C as described (92). They were treated with an additional fresh 20 mM formaldehyde and 10 mM sodium cyanoborohydride for 2 h the following day at 37 °C and the reaction was quenched with 100 mM Tris for 1 h at 37 °C. 500 μg of each channel was combined for buffer exchange on a 3 kDa molecular weight cut-off column (EMD Millipore) into 100 mM ammonium bicarbonate and digested overnight at 37 °C with mass spectrometry grade trypsin gold (Promega) at a 1:50 trypsin:protein ratio. Peptides were eluted via centrifugation and 30 µg of this digest was reserved for pre-TAILS analysis. The remaining peptides underwent enrichment using hyperbranched polyglycerol-aldehyde polymers (HPG-ALD, Flintbox, https://www.flintbox.com/public/project/1948/) at a 5:1 polymer:protein ratio. HPG-ALD binds to unblocked (i.e. trypsin-generated) amino acid termini, excluding them from the sample and enriches for blocked/labeled N-termini (93). The peptides were filtered through a 10 kDa MWCO column (EMD Millipore) to remove the polymer and obtain the TAILS fraction. TAILS and pre-TAILS fractions were desalted on a C18 Sep-Pak column (Waters) and eluted in 60:40 ACN: 1% trifluoroacetic acid. Samples were vacuum-centrifuged until dry and resuspended in 1% acetic acid for mass spectrometry.

### Mass spectrometry

Samples were analyzed on a Thermo Ultimate 3000 UHPLC interfaced with a ThermoFisher Scientific Fusion Lumos tribrid mass spectrometer system. The HPLC column was a Dionex 15 cm x 75 μm internal diameter Acclaim Pepmap C18, 2 μm, 100 Å reversed-phase capillary chromatography column. 5 μL volumes of the extract were injected and the peptides eluted from the column by an acetonitrile/0.1% formic acid gradient at a flow rate of 0.3 μL/min were introduced into the source of the mass spectrometer on-line over a 120 min gradient. The nanospray ion source is operated at 1.9 kV. The digest was analyzed using a data-dependent method with 35% collision-induced dissociation fragmentation of the most abundant peptides every 3 s and an isolation window of 0.7 m/z for ion-trap MS/MS. Scans were conducted at a maximum resolution of 120,000 for full MS. Dynamic exclusion was enabled with a repeat count of 1 and ions within 10 ppm of the fragmented mass were excluded for 60 s.

### Proteomics data analysis

Peptides were identified using a precursor mass tolerance of 10 ppm, and fragment mass tolerance of 0.6 Da. The only static modification was carbamidomethyl (C), whereas dynamic modifications included the light (28.03 Da) dimethyl formaldehyde (N-terminal, K), the heavy (34.06) dimethyl formaldehyde (N-terminal, K), oxidation (M, P), deamidation (N), acetylation (N-terminal), and Gln to pyro-Glu N-terminal cyclization. Peptides were validated using a false discovery rate (FDR) of 1% against a decoy database. Only high-confidence proteins (containing peptides at a 99% confidence level or higher) were recorded from each sample for data analysis. Pre-TAILS data required a minimum of two high-confidence peptides for protein identification and TAILS required a single peptide. Internal peptides were identified based on the criteria of having an N-terminal modification, and a sequence that does not begin prior to the third amino acid in the protein or immediately following a known signal, transit, or propeptide sequence. Peptides that met these criteria were further analyzed based on the average fold-change ratio (ADAMTS6 WT/ EA abundance) between the three technical replicate pairs. The internal peptide abundance was divided by the total protein abundance fold-change to account for differences in protein levels between groups. Peptides that met these criteria and contained a weighted ratio (ADAMTS6 WT/EA) greater than 1 underwent a t-test for significance.

### Statistics

Photomicrographs are representative data of three independent, biological replicates unless otherwise indicated. The two-tailed, unpaired Student’s *t*-test was used to obtain p values. Asterisks indicate differences with statistical significance as follows: *p≤0.05; **p≤0.01; ***p≤0.001. In the dimethyl-TAILS experiments a two-tailed, paired Student’s *t*-test was used to obtain p values. Asterisks indicate differences with statistical significance as follows: *p≤0.01, #p≤0.05.

## ACKNOWLEDGMENTS

This work was supported by the Allen Distinguished Investigator Program, through support made by The Paul G. Allen Frontiers Group and the American Heart Association (to SSA), a post-doctoral fellowship from the National Institutes of Health (F32AR063548 to TJM) and the David and Lindsay Morgenthaler Postdoctoral Fellowship (to TJM). DPR was supported by the Canadian Institutes of Heatlh Research (PJT-162099) and the Marfan Foundation (USA). The Wellcome Centre for Cell-Matrix Research is supported by funding from Wellcome (203128/Z/16/Z). CB gratefully acknowledges BBSRC funding (Ref: BB/R008221/1). We thank Dr. Francesco Ramirez for *Fbn1* and *Fbn2* mutant mice.

## Author contributions

TJM and SSA designed the study. TJM, LWW and DRM conducted experiments and acquired and analyzed the data. DPR, SAC and CB provided reagents and edited the manuscript. TJM and SSA wrote the paper.

---

**Figure 1 – figure supplement 1.**
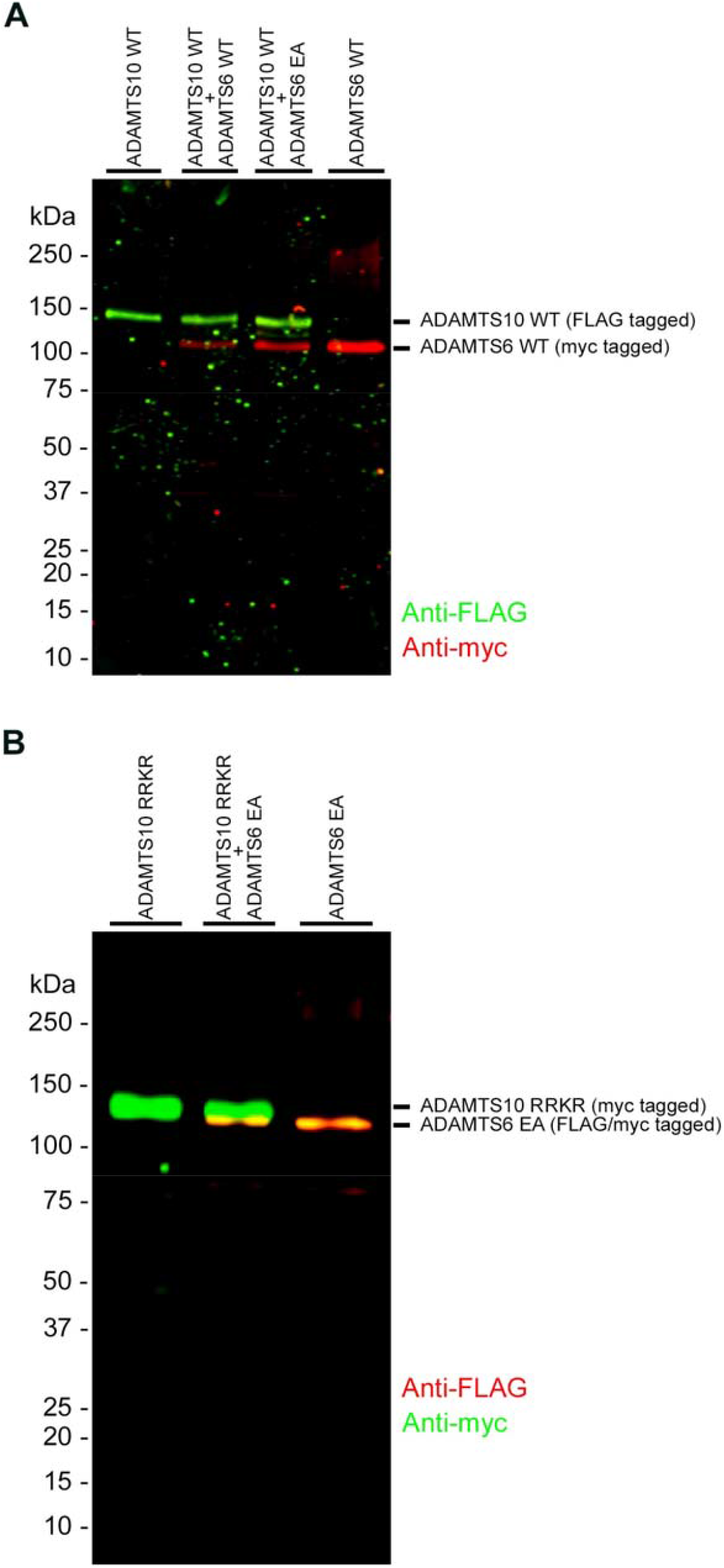
ADAMTS6 and ADAMTS10 do not cleave each other. **(A)** ADAMTS10 was co-transfected with ADAMTS6 or ADAMTS6 EA mutant. **(B)** Furin-site optimized ADAMTS10 (ADAMTS10-RRKR) was co-transfected with ADAMTS6 EA. No cleavage products of either ADAMTS6 EA or ADAMTS10 were identified.

**Figure 1 – figure supplement 2.**
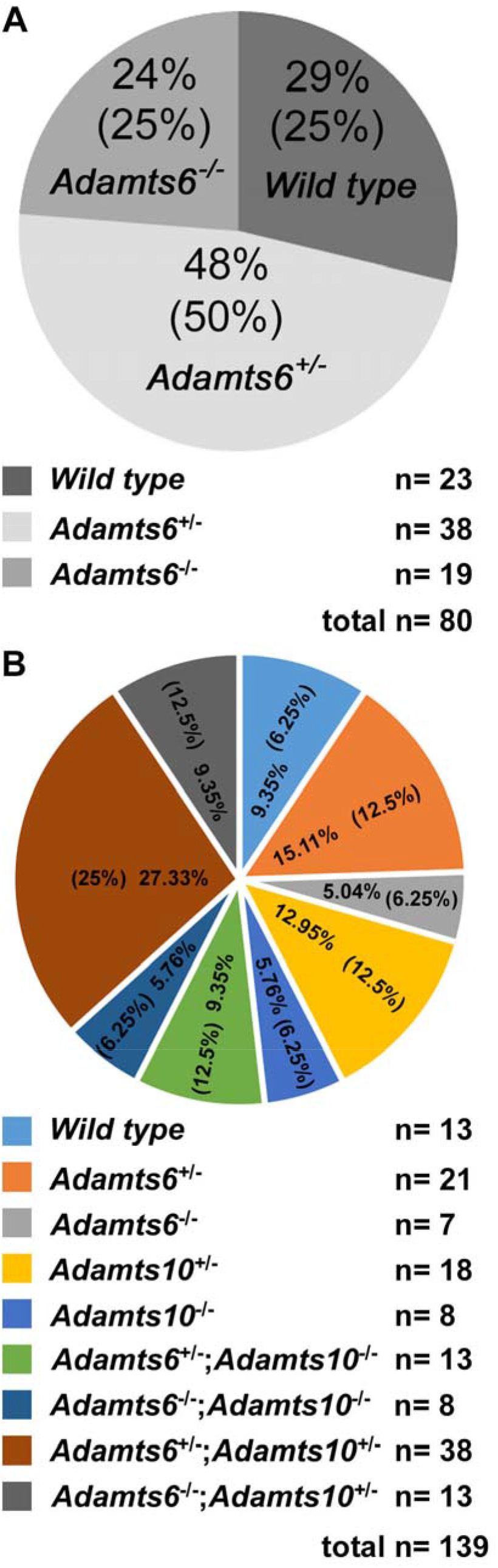
Expected Mendelian ratios are observed in mouse crosses. Mendelian ratios obtained from *Adamts6*^+/-^ intercrosses **(A)** and *Adamts6*^+/-^ *;Adamts10*^+/-^ intercrosses at E18.5 **(B)** are shown. Observed and expected (in parentheses) genotype percentages are shown in the pie-charts. The actual number of mice used in the analysis is illustrated below the pie-charts.

**Figure 1 – figure supplement 3.**
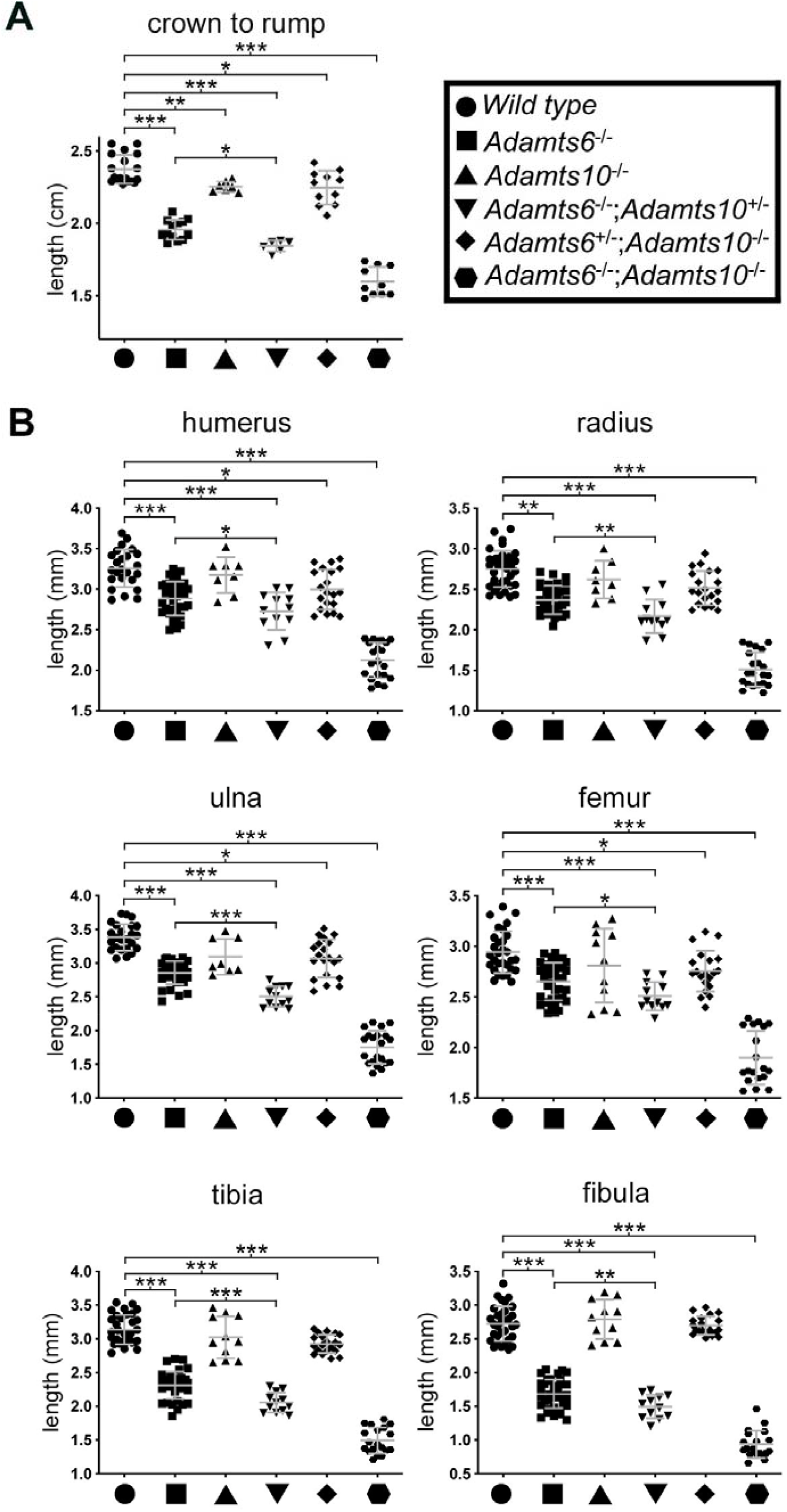
Impaired growth and shorter limb skeletal elements in combined *Adamts6* and *Adamts10* alleles. **(A)** E18.5 *Adamts6*- and *Adamts10*-deficient embryos and embryos with various combinations of these alleles have reduced crown-rump length as compared to wild type embryos. **(B)** At E18.5, embryos with various combinations of the *Adamts6*-mutant allele have shorter long bones than wild type, as shown. *Adamts10*^-/-^ long bones were not significantly shorter than those of wild type littermates. Crown-rump length, n≥ 6; bone length, n≥ 8. *p ≤0.05; **p ≤0.01; ***p ≤0.001.

**Figure 1 – figure supplement 4.**
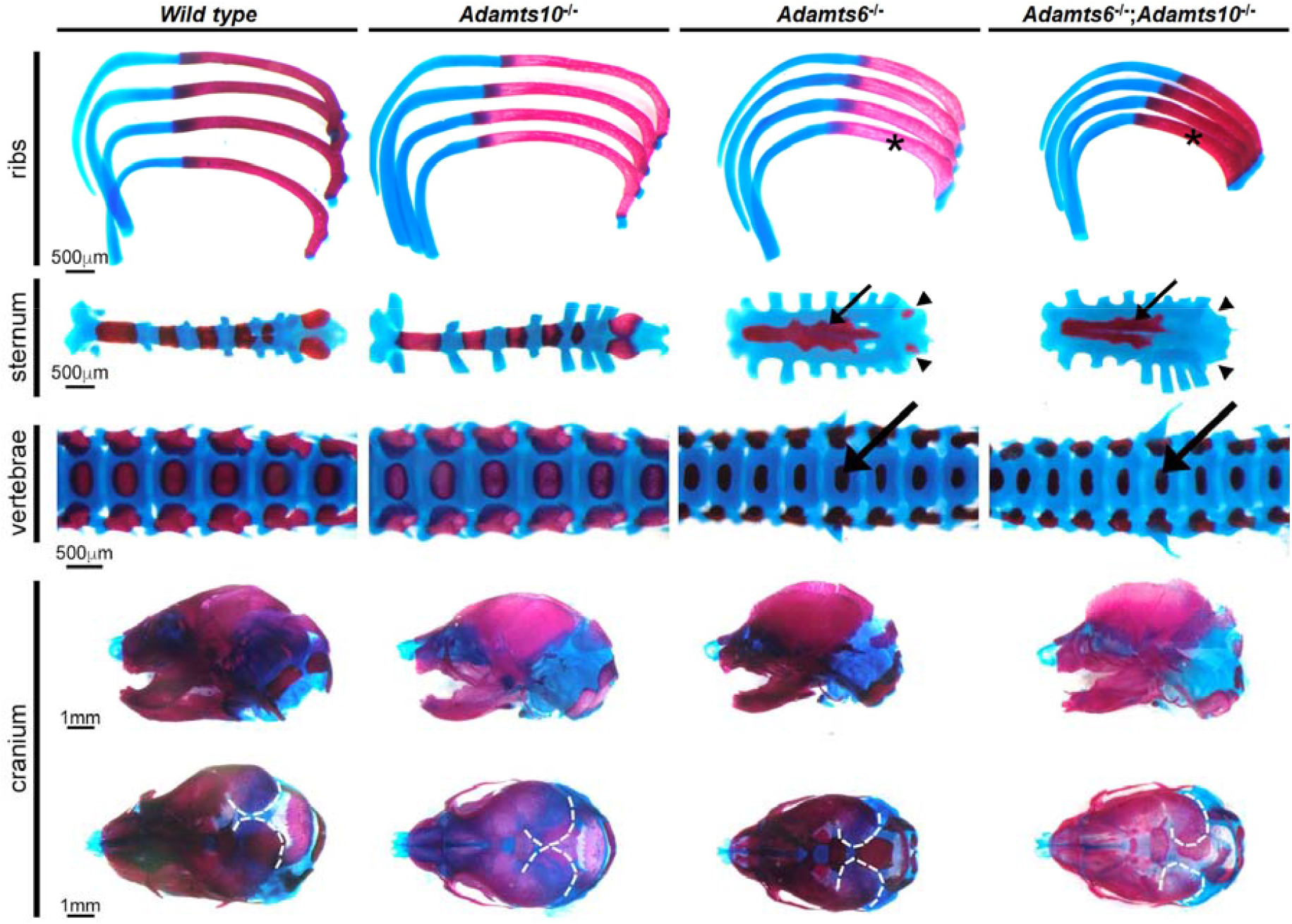
*Adamts6*-deficient embryos have a severely underdeveloped axial skeleton. Alcian blue- and alizarin red-stained E18.5 axial skeleton and craniofacial skeletal preparations show that *Adamts6^-^*^/-^ and *Adamts6^-^*^/-^;*Adamts10*^-/-^ embryos have short, stout ribs (asterisks), short and disorganized manubrium and sternum (small arrows) with under-ossified/unossified xiphoid process (arrowheads), smaller vertebral bodies (large arrows), and a small cranium with delayed mineralization of parietal bones (dashed white lines).

**Figure 1 – figure supplement 5.**
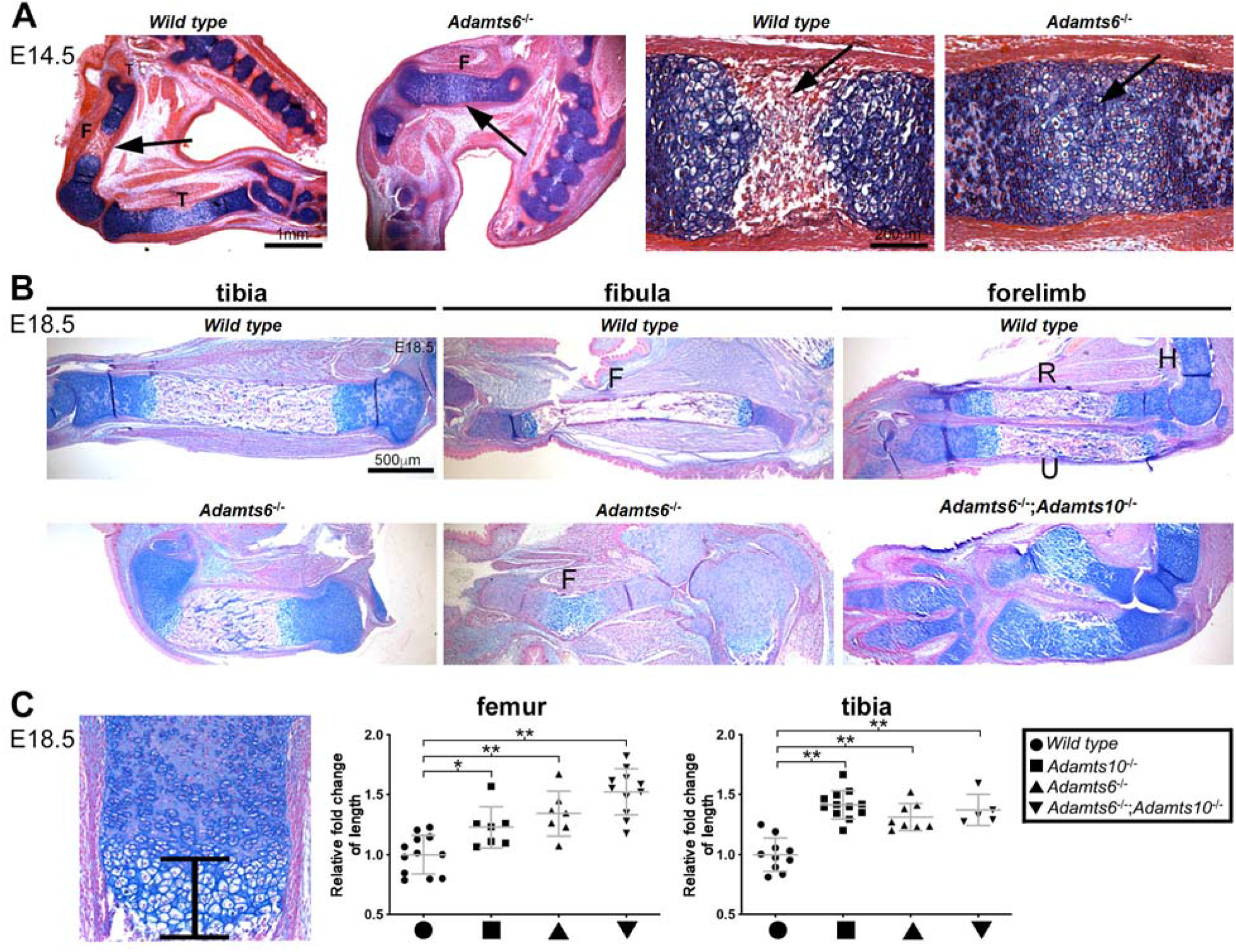
Delayed endochondral ossification, bent chondroepiphysis and impaired long bone growth in *Adamts6*-deficent hindlimbs. **(A)** Alcian blue-stained E14.5 hindlimb sections reveal delayed ossification. The arrow points to bone in wild type and hypertrophic chondrocytes in *Adamts6*^-/-^ femur, respectively. F, femur; T, tibia. (**B**) Alcian blue-stained E18.5 sections show deformed *Adamts6*^-/-^ tibia (left) and fibula (middle) as compared to wild type. Malformed radius and ulna of *Adamts6*^-/-^;*Adamts10*^-/-^ forelimb as compared to control (right). F, fibula; R, radius; U, ulna; H, humerus. (**C**) Quantification of hypertrophic chondrocyte zone in the distal femur and proximal tibia shows increased thickness in the mutants. n≥ 6. *p ≤0.05; **p ≤0.01.

**Figure 1 – figure supplement 6.**
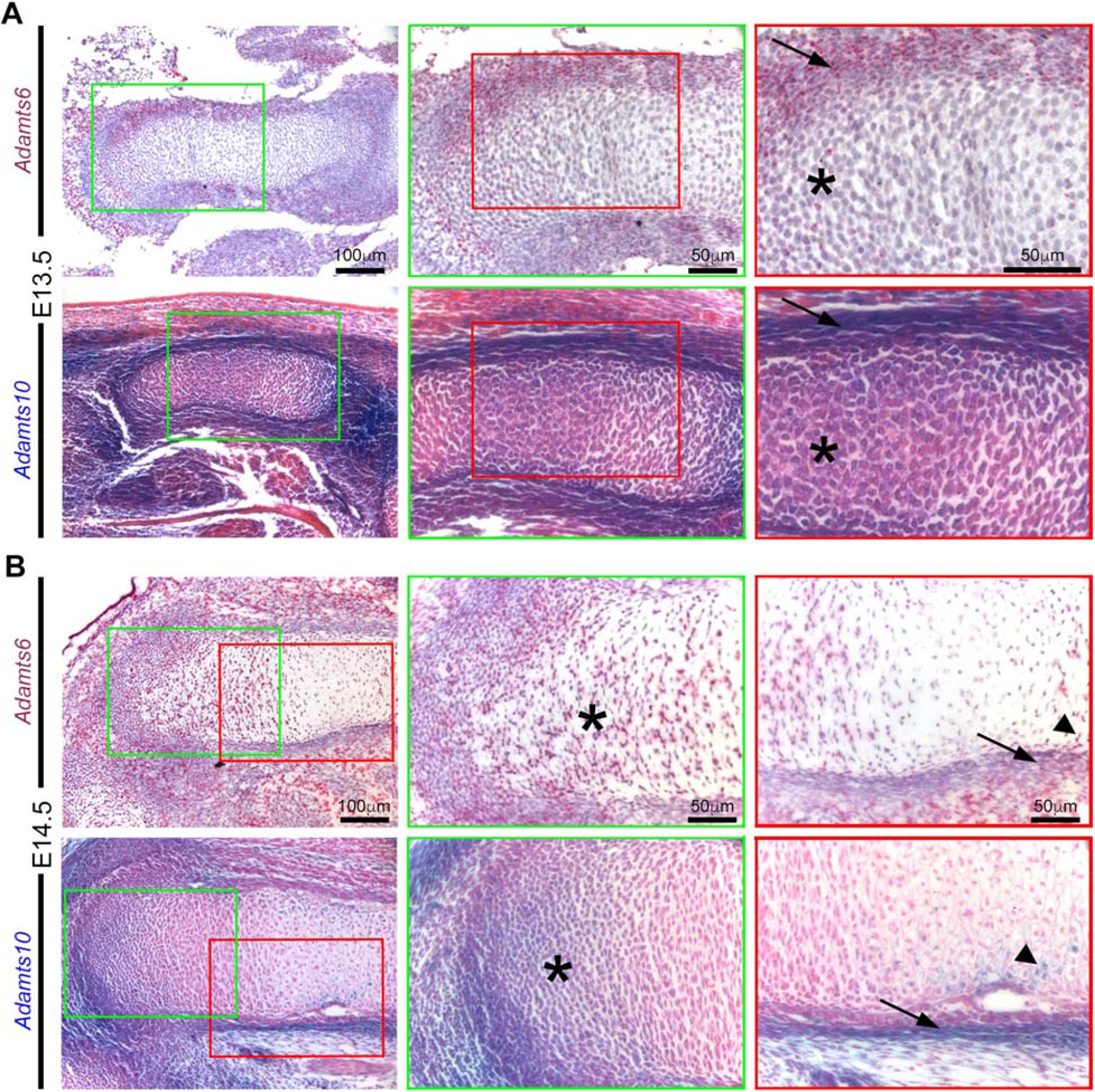
*Adamts6* and *Adamts10* mRNAs show overlapping expression in the developing hindimbs. *Adamts6* (RNA in situ hybridization (red signal)) and *Adamts10* (β-gal staining of *Adamts10*^+/-^ tissues, blue nuclei) are expressed in E13.5 **(A)** and E14.5 **(B)** perichondrium (arrows), resting chondrocytes (asterisks) and peripheral hypertrophic chondrocytes at the site of vascular invasion that form the primary ossific centers (arrowheads).

**Figure 2 – figure supplement 1.**
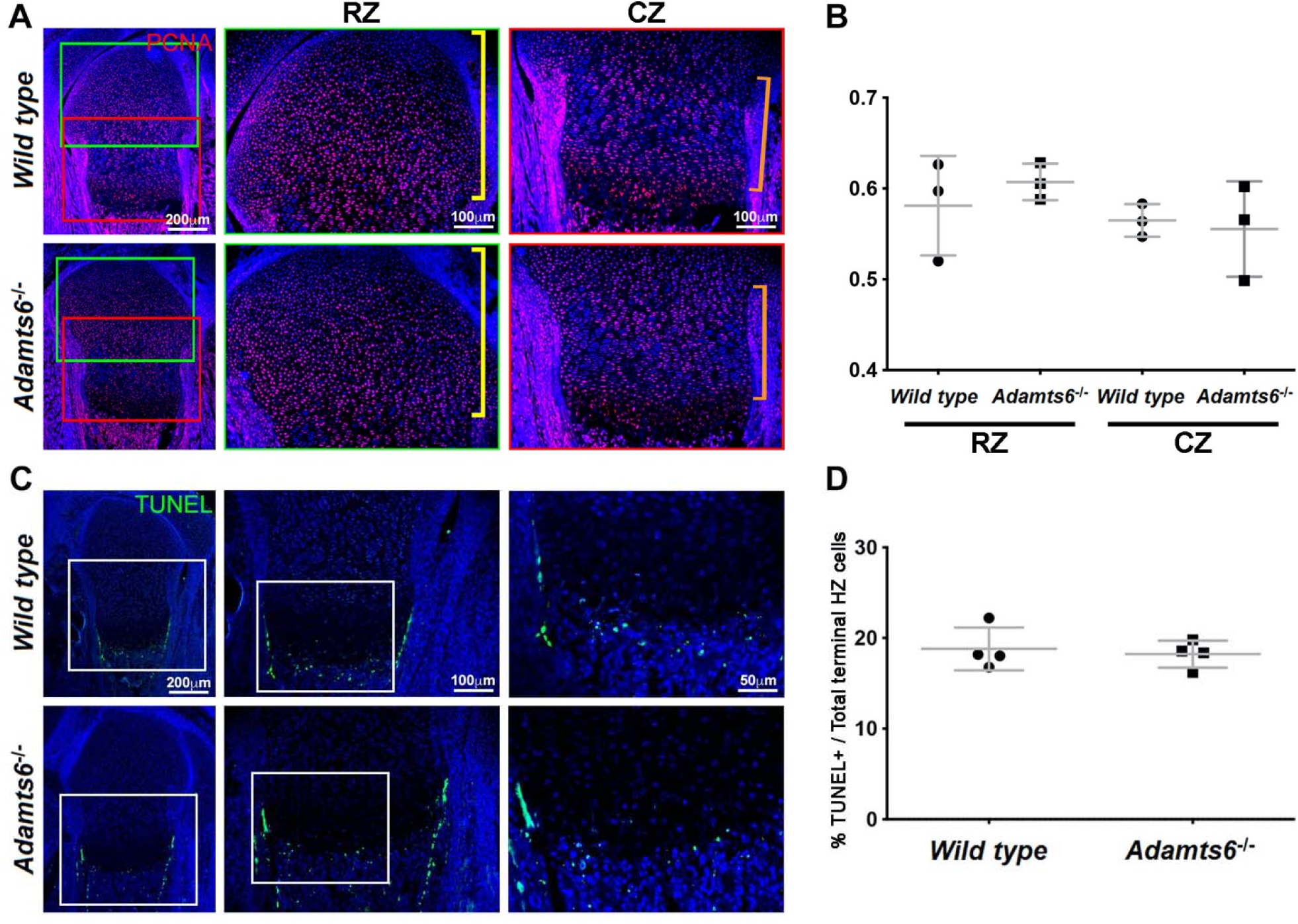
No change in proliferation or cell death in *Adamts6*^-/-^ cartilage. **(A-B)** PCNA+ nuclei (red) **(A)** in E18.5 distal femur were quantified in the resting zone (RZ; yellow brackets) and columnar zone (CZ; orange brackets and quantified **(B)**. N=4. (**C-D)** TUNEL+ cells (green) (**C**) in E18.5 proximal femur were counted and quantified **(D)** at the chondro-osseous junction (terminal hypertrophic chondrocytes).

**Figure 3 – figure supplement 1.**
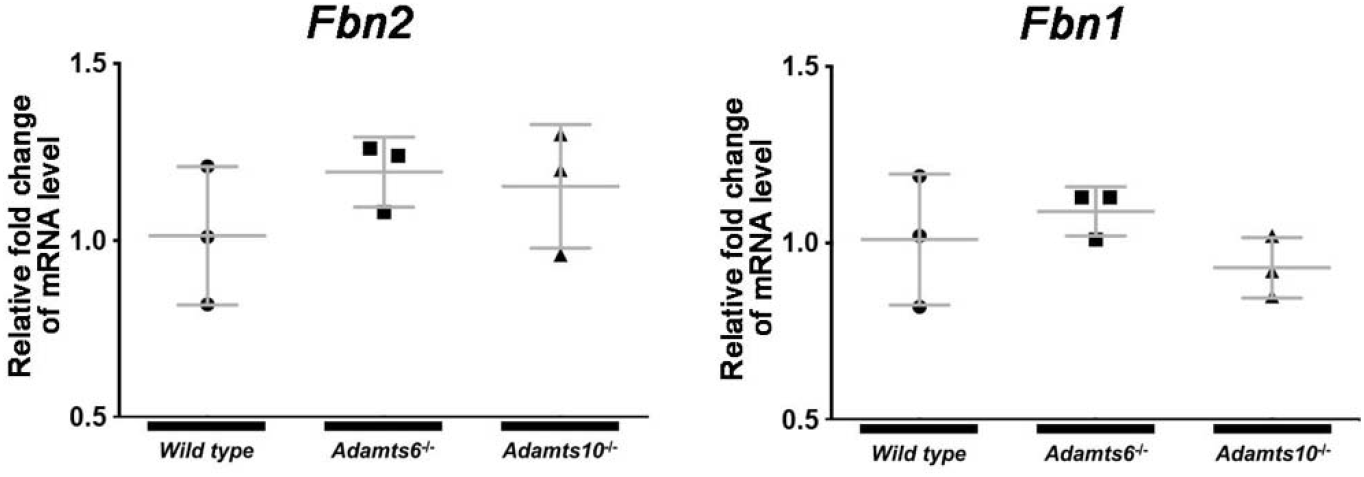
Comparable *Fbn2* and *Fbn1* mRNA levels in *Adamts6 Adamts10* mutant hindlimbs. qRT-PCR was done in E18.5 hindlimbs. n=3.

**Figure 4 – figure supplement 1.**
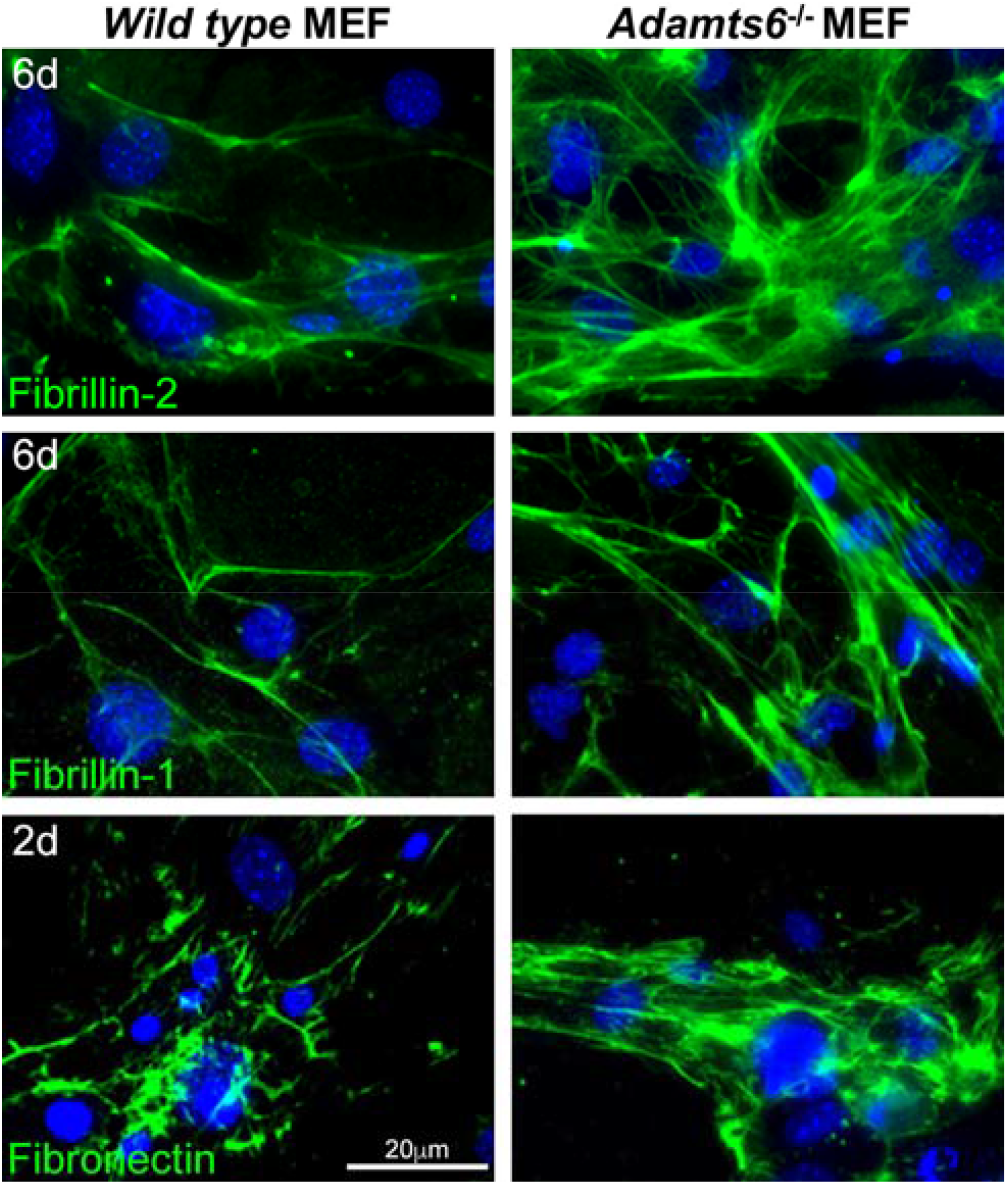
Increased fibrillin-1, fibrillin-2 and fibronectin microfibril staining in the absence of *ADAMTS6*. 6-day cultures of *Adamts6*^-/-^ mouse embryo fibroblasts (MEFs) have increased fibrillin-2 and fibrillin-1 staining compared to wild type MEFs. Fibronectin staining is similarly increased in day 2 cultures. Nuclei are stained with DAPI (blue).

**Figure 5 – figure supplement 1.**
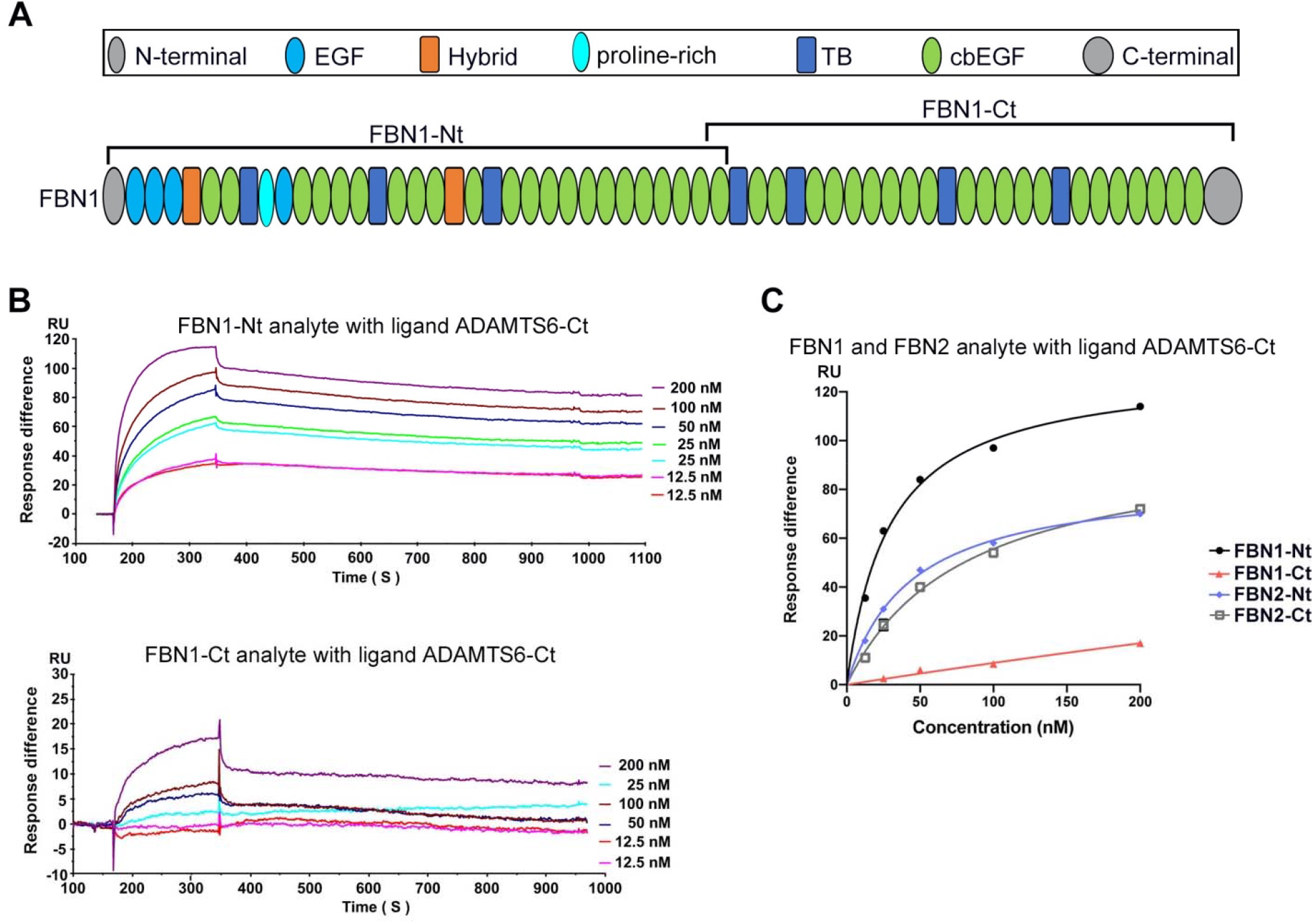
ADAMTS6 binds directly to fibrillin-1. **(A)** Domain structure of fibrillin-1 recombinant constructs. **(B-C)** Biacore analysis shows that fibrillin-2-Nt and fibrillin-2-Ct recombinant fragments bind strongly to ADAMTS6-Ct, with FBN1-Nt binding the strongest among FBN1-Nt, FBN1-Ct, F BN2-Nt and FBN2-Ct.

**Figure 6 – figure supplement 1.**
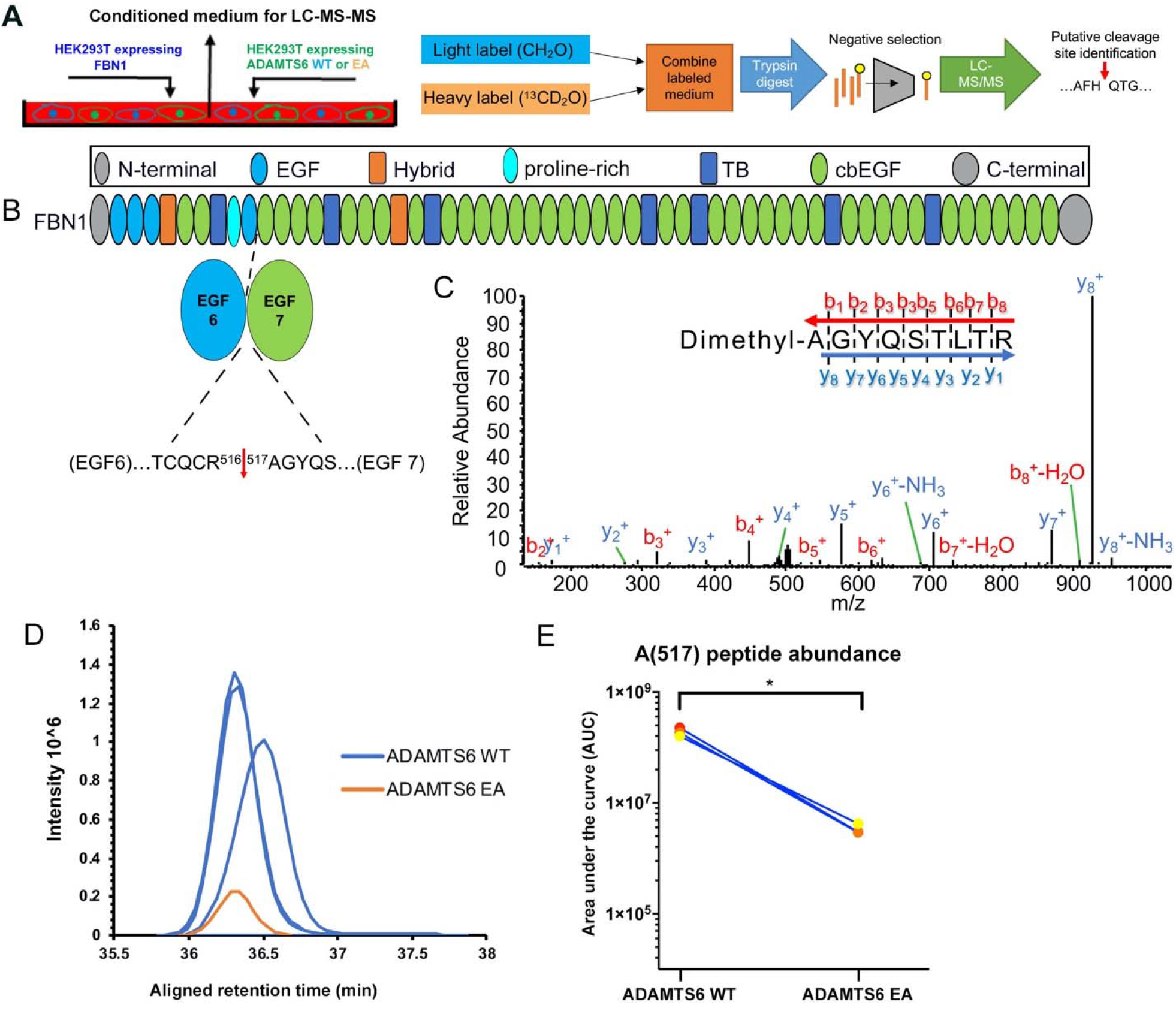
ADAMTS6 cleaves fibrillin-1. **(A)** Schematic of the experimental approach. Proteins from conditioned medium of co-cultured HEK293F cells stably expressing fibrillin-1 and cells expressing either ADAMTS6 WT or ADAMTS6 EA (inactive) were labeled by reductive dimethylation using stable formaldehyde isotopes and analyzed by LC-MS/MS in the TAILS workflow described in detail in the Methods section. **(B)** Schematic of the fibrillin-1 domain structure indicating the location of the identified cleavage site. **(C)** Annotated MS2 spectrum of the fibrillin-1 peptide shows the b-(N-terminal preserved) and y-type (C-terminal preserved) ions generated via amide bond cleavage during collision-induced dissociation revealing the peptide backbone sequence. **(D)** The retention time-aligned extracted ion chromatographs show the light dimethyl labeled AGYQSLTR peptide (blue) from the ADAMTS6 WT medium compared to the isotopically heavy dimethyl labeled peptide (orange) from the ADAMTS6 EA medium. **(E)** The area under the extracted ion chromatogram was quantified in 3 TAILS replicates and plotted. Significance was determined using a two-tailed, paired Student t-test, *p ≤ 0.05.

**Figure 6 – figure supplement 2.**
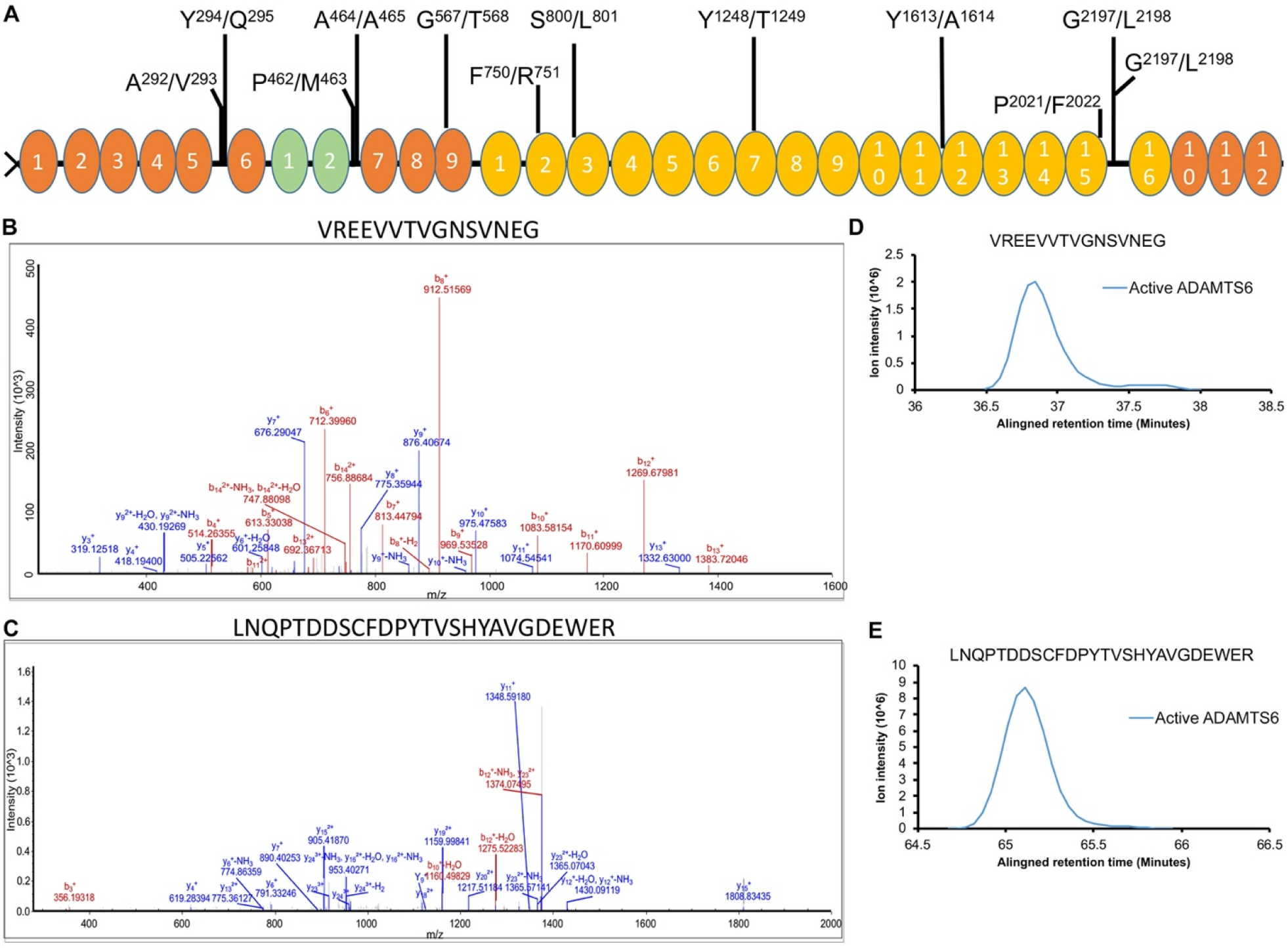
ADAMTS6 cleaves fibronectin at multiple sites. **(A)** Schematic of the domain structure of fibronectin and locations of the identified cleavage sites. **(B-C**) Annotated MS2 spectra of the VREEVVTVGNSVNEG^2197^ (**B**) and ^2198^LNQPTDDSCFDPYTVSHYAVGDEWER (**C**) peptides, each identifying a cleavage at Gly^2197^- and Leu^2198^. (**D-E**) Extracted ion chromatograms showing peptide intensity and retention time in the ADAMTS6 medium for peptides VREEVVTVGNSVNEG^2197^ (**D**) and ^2198^LNQPTDDSCFDPYTVSHYAVGDEWER (**E**). A matching chromatographic trace was not detected in the ADAMTS6 EA medium.

**Figure 7 – figure supplement 1.**
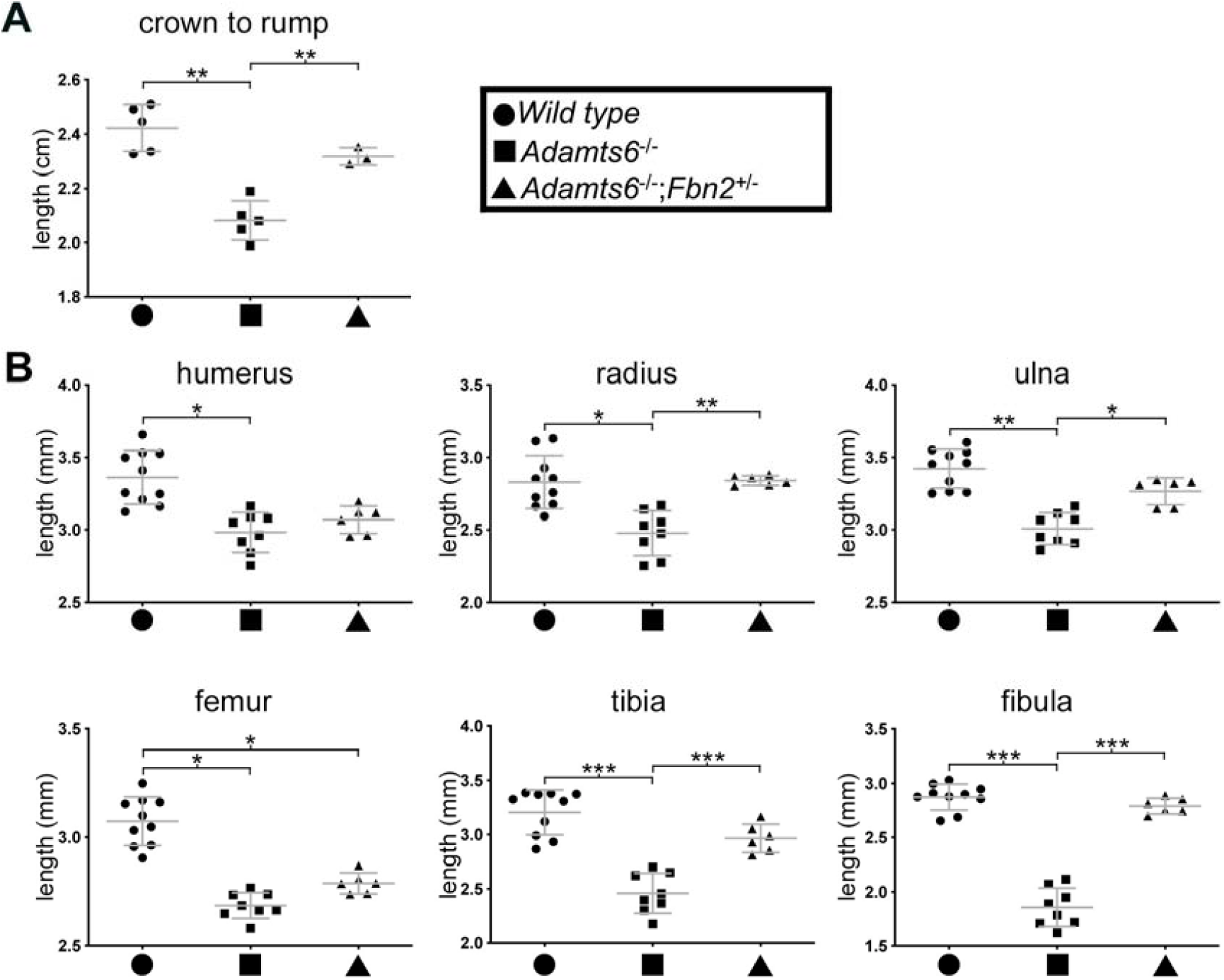
Amelioration of reduced body length and long bone shortening in *Adamts6*-deficient embryos by *Fbn2* hemizygosity. Reduced crown-rump length **(A)** and reduced radius, ulna, tibia and fibula length **(B)** of *Adamts6^-^*^/-^ E18.5 embryos are ameliorated by *Fbn2* heterozygosity. Crown-rump length, n≥ 3; bone length, n≥ 6. *p ≤0.05; **p ≤0.01; ***p ≤0.001.

**Figure 7 – figure supplement 2.**
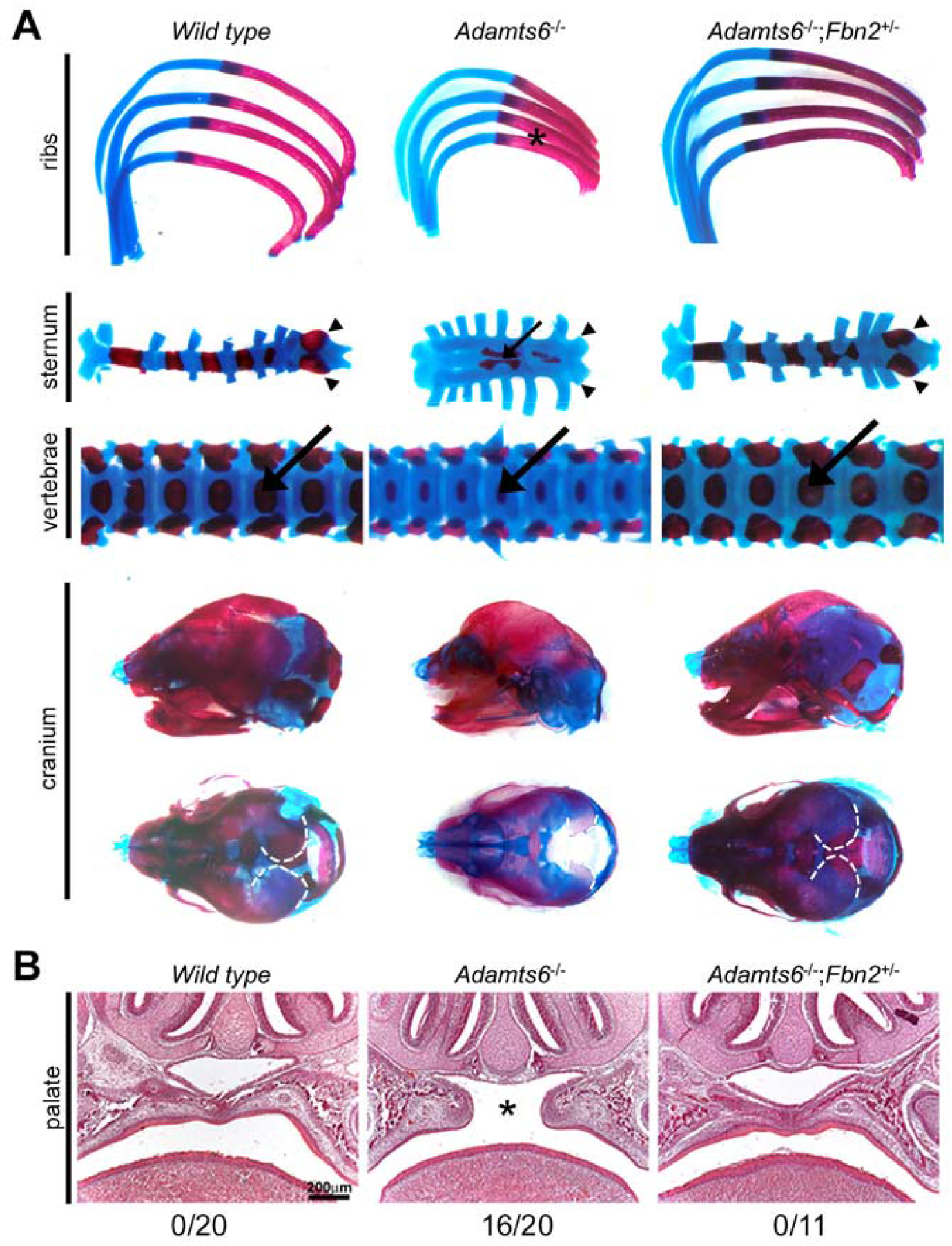
*Fbn2* haploinsufficiency reverses axial skeleton and craniofacial anomalies in *Adamts6^-^*^/-^ mice. The images illustrate restoration of sternal segmental ossific centers (arrows) and restoration of length, ossification and relative proportions of the ribs, sternal segments and xiphoid process (arrowheads), vertebral bodies (thick arrows) and appendages, and restoration of skull dimensions and parietal ossification (dashed white line). **(B)** Hematoxylin and eosin-stained coronal sections from E18.5 embryos show that *Fbn2* haploinsufficiency reverses cleft secondary palate observed in the majority of *Adamts6*^-/-^ mutants (asterisk; incidence listed below the respective panels).

**Figure 7 – figure supplement 3.**
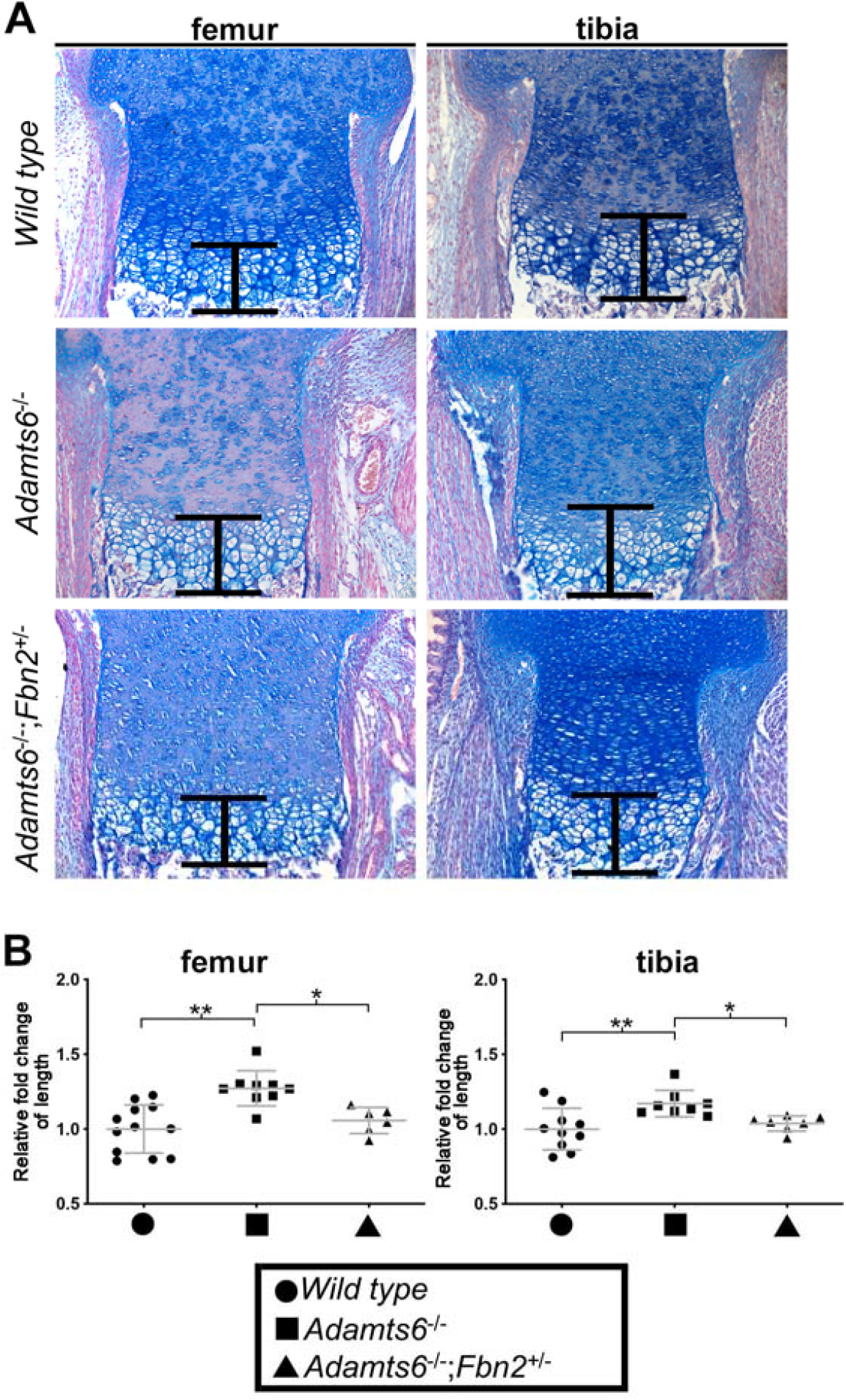
*Adamts6*-deficient femoral hypertrophic zone (HZ) thickening is reversed by *Fbn2* haploinsufficiency. (**A)** Alcian blue-stained section with the HZ marked by the bracket. **(B)** Quantitation of the relative thickness of the HZ in the indicated genotypes. n≥ 6. *p ≤0.05; **p ≤0.01.

**Figure 7 – figure supplement 4.**
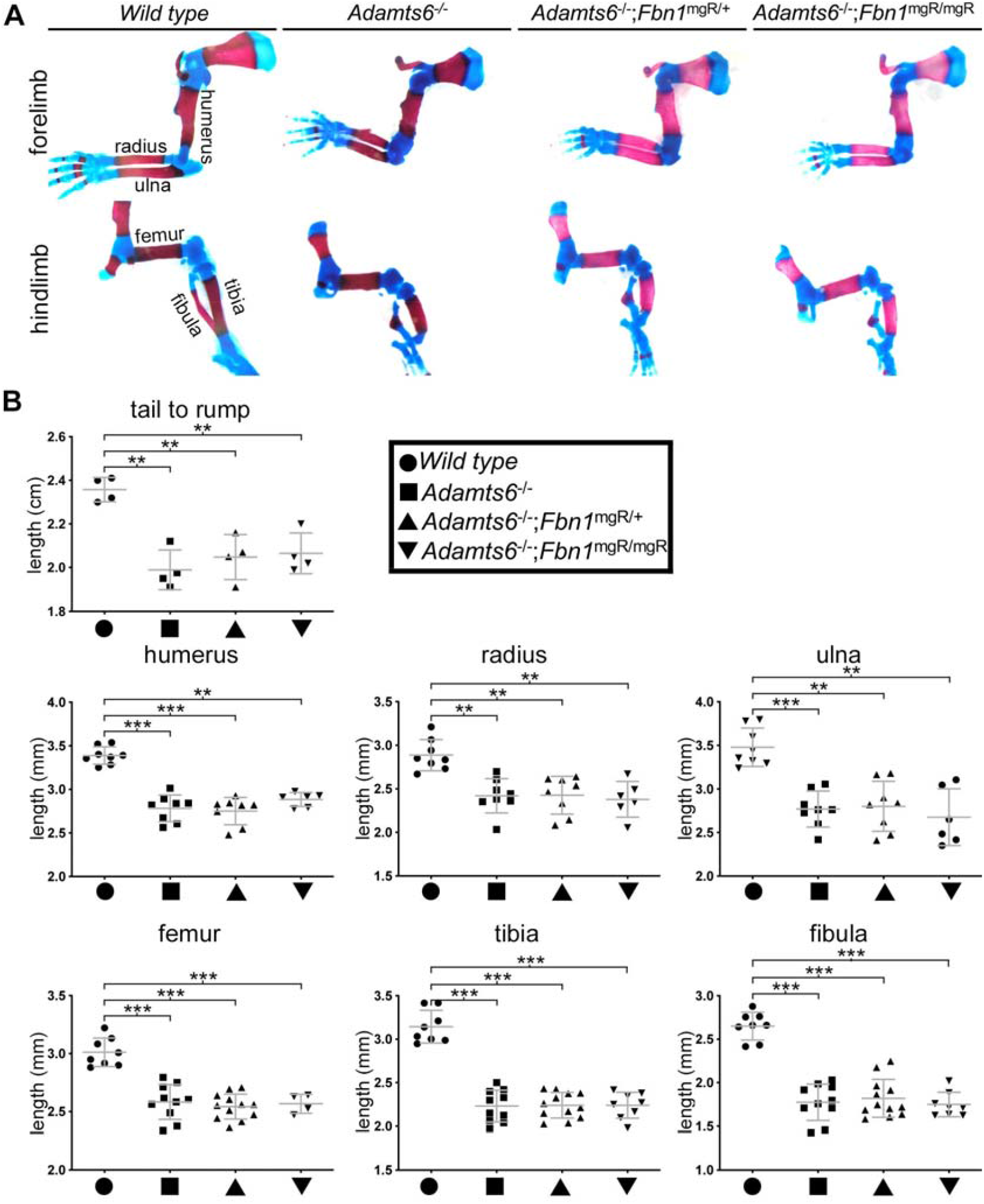
No improvement in long bone shortening (A,B) or crown-rump length (B) of *Adamts6*^-/-^ embryos by hemizygosity or homozygosity of a *Fbn1* mutant allele, *Fbn1*^mgR^. Measurements were made in E18.5 embryos of the indicated genotypes. Crown-rump length, n≥ 4; bone length, n≥ 4. *p ≤0.05; **p ≤0.01; ***p ≤0.001.

**Figure 7 – figure supplement 5.**
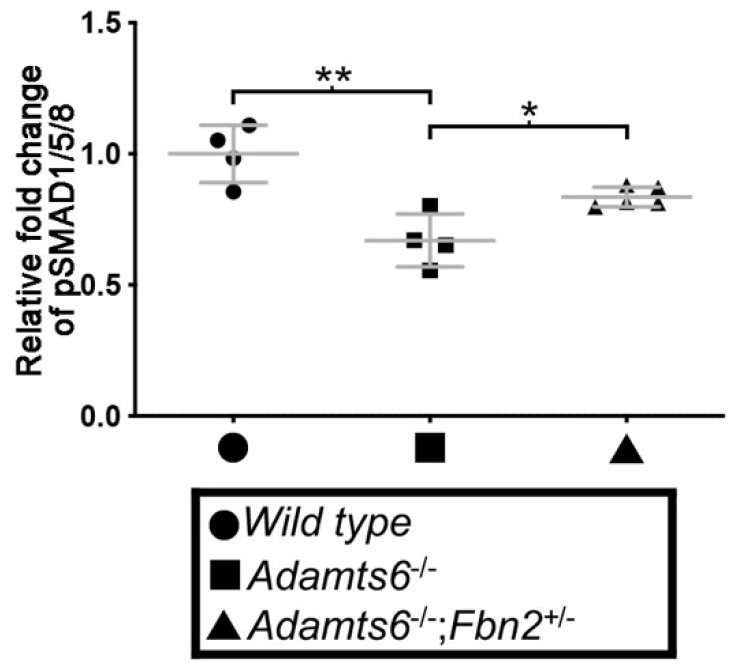
BMP signaling is reduced in *Adamts6*-deficient distal femoral cartilage and restored in *Adamts6*^-/-^;*Fbn2*^+/-^ limbs. Quantification of positive nuclei as a percentage of total nuclei identified by DAPI staining in histological sections stained with anti-pSmad1/5/8 in Figure 7E depicted as fold-change relative to wild type staining. n= 4-5. *p ≤0.05; **p ≤0.01.

**Figure 7 – figure supplement 6.**
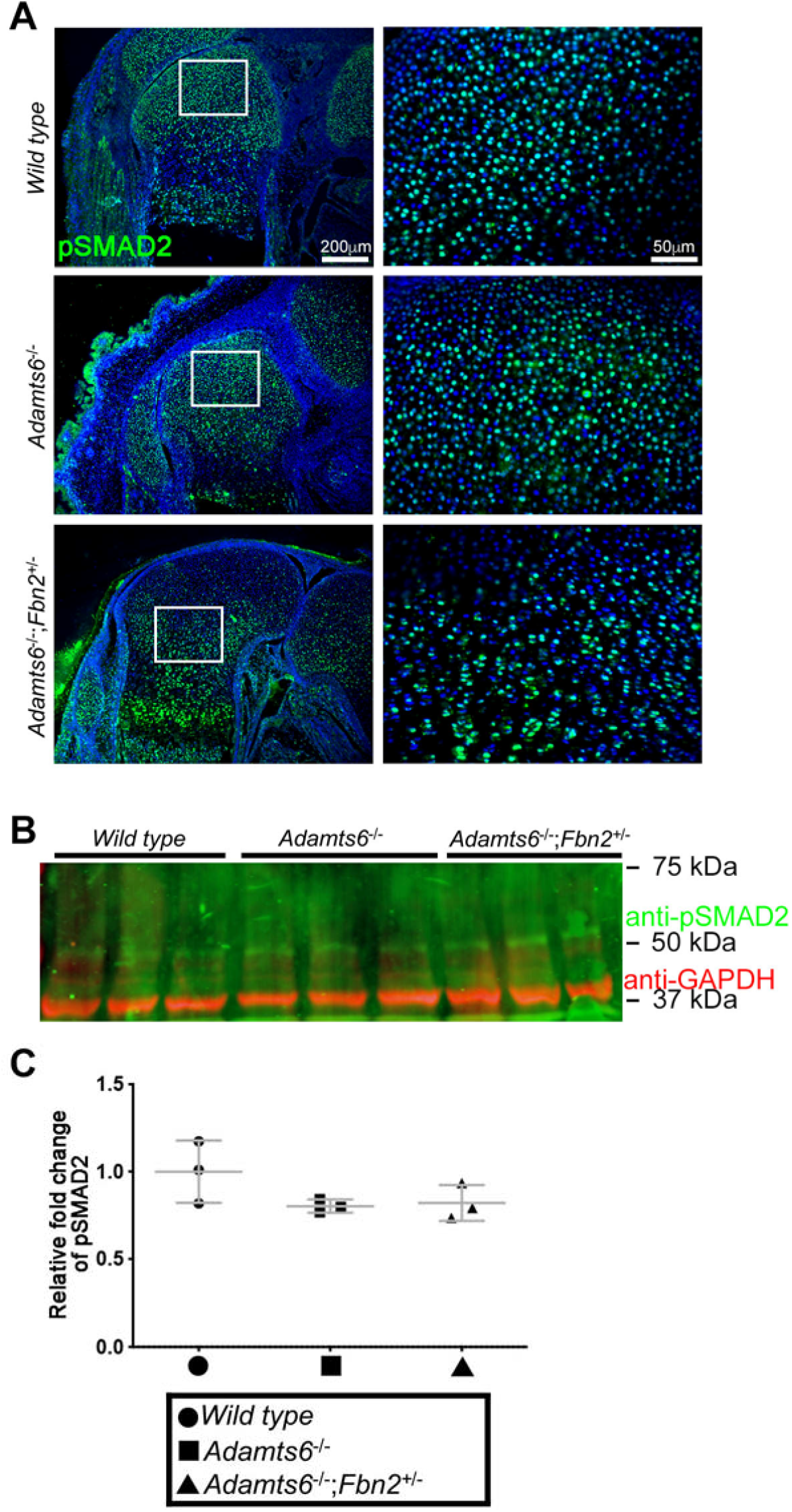
TGFβ signaling is unaffected in *Adamts6*-deficient distal femur. **(A)** No change in pSMAD2 staining in *Adamts6*^-/-^ femur as compared to wild type or *Adamts6*^-/-^;*Fbn2*^+/-^ femur. Western blot analysis shows no change in anti-pSmad2 (green; 52 kDa) content in *Adamts6*-deficient E18.5 hindlimb lysates. Anti-GAPDH (red, 37 kDa) was used as a loading control. **(C)** Quantification of anti-pSmad2 signal in (A) using GAPDH signal intensity as the control. n= 3. p* ≤0.05.

**Supplemental Table 1.** Antibodies.

| **Name** | **Source** | **dilution** |
| --- | --- | --- |
| **anti-Fibrillin-2-gly** | (94) | IF: 1:300 |
| **anti-mFbn1-C** | (95) | IF: 1:500 |
| **Anti-MAGP1** | (96) | IF: 1:200 |
| **anti-Sox9** | Millipore AB5535 | IF: 1:300 |
| **anti-Acan** | Millipore AB1031 | IF: 1:400 |
| **anti-CLP** | DHSB 9/30/8-A4-c | IF: 1:100 |
| **anti-Col X** | Abcam ab58632 | IF: 1:1000 |
| **Anti-PCNA** | Cell Signaling 2586S | IF: 1:200 |
| **anti-rFBN2-C** | (21) | WB: 1:500 |
| **anti-His** | R&D MAB050 | IF: 1:400<br>WB: 1:1000 |
| **anti-pSmad5** | Abcam ab92698 | WB: 1:1000 |
| **Anti-pSmad1/5** | Cell Signaling 9516 | IF: 1:200 |
| **anti-pSmad2** | Cell Signaling 3108 | WB: 1:1000<br>IF: 1:200 |
| **anti-GAPDH** | EMD Millipore MAB374 | WB: 1:5000 |

IF, immunofluorescence; WB, western blot

**Supplemental Table 2.**
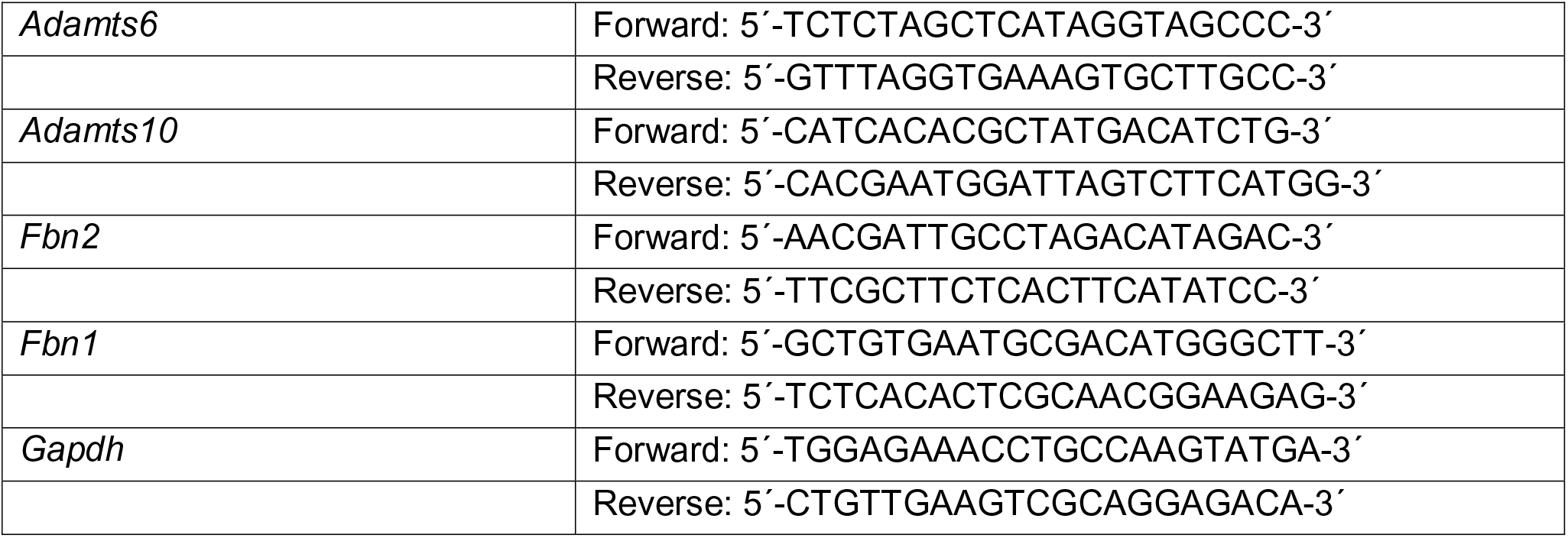
Quantitative Real-Time PCR primers.

## REFERENCES

1. Huxley-Jones J, Robertson DL, Boot-Handford RP. On the origins of the extracellular matrix in vertebrates. Matrix Biol. 2007;26(1):2–11. Epub 2006/10/24. doi: 10.1016/j.matbio.2006.09.008. PubMed PMID: 17055232.

2. Huxley-Jones J, Apte SS, Robertson DL, Boot-Handford RP. The characterisation of six ADAMTS proteases in the basal chordate Ciona intestinalis provides new insights into the vertebrate ADAMTS family. Int J Biochem Cell Biol. 2005;37(9):1838–45. PubMed PMID: 15899586.

3. Huxley-Jones J, Clarke TK, Beck C, Toubaris G, Robertson DL, Boot-Handford RP. The evolution of the vertebrate metzincins; insights from Ciona intestinalis and Danio rerio. BMC Evol Biol. 2007;7:63. Epub 2007/04/19. doi: 10.1186/1471-2148-7-63. PubMed PMID: 17439641; PMCID: PMC1867822.

4. Dubail J, Apte SS. Insights on ADAMTS proteases and ADAMTS-like proteins from mammalian genetics. Matrix Biol. 2015;44–46:24-37. doi: 10.1016/j.matbio.2015.03.001. PubMed PMID: 25770910.

5. Mead TJ, Apte SS. ADAMTS proteins in human disorders. Matrix Biol. 2018;71–72:225-39. Epub 2018/06/10. doi: 10.1016/j.matbio.2018.06.002. PubMed PMID: 29885460; PMCID: PMC6146047.

6. Ramirez F, Caescu C, Wondimu E, Galatioto J. Marfan syndrome; A connective tissue disease at the crossroads of mechanotransduction, TGFβ signaling and cell stemness. Matrix Biol. 2018;71–72:82-9. Epub 2017/08/08. doi: 10.1016/j.matbio.2017.07.004. PubMed PMID: 28782645; PMCID: PMC5797509.

7. Thomson J, Singh M, Eckersley A, Cain SA, Sherratt MJ, Baldock C. Fibrillin microfibrils and elastic fibre proteins: Functional interactions and extracellular regulation of growth factors. Semin Cell Dev Biol. 2019;89:109–17. Epub 2018/07/18. doi: 10.1016/j.semcdb.2018.07.016. PubMed PMID: 30016650; PMCID: PMC6461133.

8. Ricard-Blum S, Vallet SD. Fragments generated upon extracellular matrix remodeling: Biological regulators and potential drugs. Matrix Biol. 2019;75–76:170-89. Epub 2017/11/15. doi: 10.1016/j.matbio.2017.11.005. PubMed PMID: 29133183.

9. Shin SJ, Yanagisawa H. Recent updates on the molecular network of elastic fiber formation. Essays Biochem. 2019;63(3):365–76. Epub 2019/08/10. doi: 10.1042/ebc20180052. PubMed PMID: 31395654.

10. Kozel BA, Mecham RP. Elastic fiber ultrastructure and assembly. Matrix Biol. 2019;84:31–40. Epub 2019/11/02. doi: 10.1016/j.matbio.2019.10.002. PubMed PMID: 31669522.

11. Sabatier L, Miosge N, Hubmacher D, Lin G, Davis EC, Reinhardt DP. Fibrillin-3 expression in human development. Matrix Biol. 2011;30(1):43–52. Epub 2010/10/26. doi: S0945-053X(10)00153-8 [pii] 10.1016/j.matbio.2010.10.003. PubMed PMID: 20970500.

12. Zhang H, Apfelroth SD, Hu W, Davis EC, Sanguineti C, Bonadio J, Mecham RP, Ramirez F. Structure and expression of fibrillin-2, a novel microfibrillar component preferentially located in elastic matrices. J Cell Biol. 1994;124(5):855–63. Epub 1994/03/01. PubMed PMID: 8120105; PMCID: PMC2119952.

13. Zhang H, Hu W, Ramirez F. Developmental expression of fibrillin genes suggests heterogeneity of extracellular microfibrils. J Cell Biol. 1995;129(4):1165–76. PubMed PMID: 7744963; PMCID: 2120487.

14. Corson GM, Charbonneau NL, Keene DR, Sakai LY. Differential expression of fibrillin-3 adds to microfibril variety in human and avian, but not rodent, connective tissues. Genomics. 2004;83(3):461–72. Epub 2004/02/14. doi: 10.1016/j.ygeno.2003.08.023 S0888754303002738 [pii]. PubMed PMID: 14962672.

15. Yip RKH, Chan D, Cheah KSE. Mechanistic insights into skeletal development gained from genetic disorders. Curr Top Dev Biol. 2019;133:343–85. Epub 2019/03/25. doi: 10.1016/bs.ctdb.2019.02.002. PubMed PMID: 30902258.

16. Robinson PN, Arteaga-Solis E, Baldock C, Collod-Beroud G, Booms P, De Paepe A, Dietz HC, Guo G, Handford PA, Judge DP, Kielty CM, Loeys B, Milewicz DM, Ney A, Ramirez F, Reinhardt DP, Tiedemann K, Whiteman P, Godfrey M. The molecular genetics of Marfan syndrome and related disorders. J Med Genet. 2006;43(10):769–87. Epub 2006/03/31. doi: jmg.2005.039669 [pii] 10.1136/jmg.2005.039669. PubMed PMID: 16571647; PMCID: 2563177.

17. Arteaga-Solis E, Gayraud B, Lee SY, Shum L, Sakai L, Ramirez F. Regulation of limb patterning by extracellular microfibrils. J Cell Biol. 2001;154(2):275–81. PubMed PMID: 11470817.

18. Sengle G, Carlberg V, Tufa SF, Charbonneau NL, Smaldone S, Carlson EJ, Ramirez F, Keene DR, Sakai LY. Abnormal Activation of BMP Signaling Causes Myopathy in Fbn2 Null Mice. PLoS Genet. 2015;11(6):e1005340. doi: 10.1371/journal.pgen.1005340. PubMed PMID: 26114882; PMCID: PMC4482570.

19. Charbonneau NL, Dzamba BJ, Ono RN, Keene DR, Corson GM, Reinhardt DP, Sakai LY. Fibrillins can co-assemble in fibrils, but fibrillin fibril composition displays cell-specific differences. J Biol Chem. 2003;278(4):2740–9. Epub 2002/11/14. doi: 10.1074/jbc.M209201200 M209201200 [pii]. PubMed PMID: 12429739.

20. Marson A, Rock MJ, Cain SA, Freeman LJ, Morgan A, Mellody K, Shuttleworth CA, Baldock C, Kielty CM. Homotypic fibrillin-1 interactions in microfibril assembly. J Biol Chem. 2005;280(6):5013–21. Epub 2004/12/01. doi: M409029200 [pii] 10.1074/jbc.M409029200. PubMed PMID: 15569675.

21. Lin G, Tiedemann K, Vollbrandt T, Peters H, Batge B, Brinckmann J, Reinhardt DP. Homo- and heterotypic fibrillin-1 and −2 interactions constitute the basis for the assembly of microfibrils. J Biol Chem. 2002;277(52):50795–804. Epub 2002/10/26. doi: 10.1074/jbc.M210611200 M210611200 [pii]. PubMed PMID: 12399449.

22. Nistala H, Lee-Arteaga S, Smaldone S, Siciliano G, Carta L, Ono RN, Sengle G, Arteaga-Solis E, Levasseur R, Ducy P, Sakai LY, Karsenty G, Ramirez F. Fibrillin-1 and −2 differentially modulate endogenous TGF-beta and BMP bioavailability during bone formation. J Cell Biol. 2010;190(6):1107–21. Epub 2010/09/22. doi: jcb.201003089 [pii] 10.1083/jcb.201003089. PubMed PMID: 20855508.

23. Dagoneau N, Benoist-Lasselin C, Huber C, Faivre L, Megarbane A, Alswaid A, Dollfus H, Alembik Y, Munnich A, Legeai-Mallet L, Cormier-Daire V. ADAMTS10 mutations in autosomal recessive Weill-Marchesani syndrome. Am J Hum Genet. 2004;75(5):801–6. PubMed PMID: 15368195.

24. Faivre L, Gorlin RJ, Wirtz MK, Godfrey M, Dagoneau N, Samples JR, Le Merrer M, Collod-Beroud G, Boileau C, Munnich A, Cormier-Daire V. In frame fibrillin-1 gene deletion in autosomal dominant Weill-Marchesani syndrome. J Med Genet. 2003;40(1):34–6. PubMed PMID: 12525539.

25. Hubmacher D, Apte SS. ADAMTS proteins as modulators of microfibril formation and function. Matrix Biol. 2015;47:34–43. doi: 10.1016/j.matbio.2015.05.004. PubMed PMID: 25957949; PMCID: PMC4731137.

26. Karoulias SZ, Taye N, Stanley S, Hubmacher D. The ADAMTS/Fibrillin Connection: Insights into the Biological Functions of ADAMTS10 and ADAMTS17 and Their Respective Sister Proteases. Biomolecules. 2020;10(4). Epub 2020/04/16. doi: 10.3390/biom10040596. PubMed PMID: 32290605; PMCID: PMC7226509.

27. Cain SA, Mularczyk EJ, Singh M, Massam-Wu T, Kielty CM. ADAMTS-10 and −6 differentially regulate cell-cell junctions and focal adhesions. Sci Rep. 2016;6:35956. Epub 2016/10/26. doi: 10.1038/srep35956. PubMed PMID: 27779234; PMCID: PMC5078793.

28. Kutz WE, Wang LW, Bader HL, Majors AK, Iwata K, Traboulsi EI, Sakai LY, Keene DR, Apte SS. ADAMTS10 Protein Interacts with Fibrillin-1 and Promotes Its Deposition in Extracellular Matrix of Cultured Fibroblasts. J Biol Chem. 2011;286(19):17156–67. Epub 2011/03/16. doi: M111.231571 [pii] 10.1074/jbc.M111.231571. PubMed PMID: 21402694; PMCID: 3089559.

29. Mularczyk EJ, Singh M, Godwin ARF, Galli F, Humphreys N, Adamson AD, Mironov A, Cain SA, Sengle G, Boot-Handford RP, Cossu G, Kielty CM, Baldock C. ADAMTS10-mediated tissue disruption in Weill-Marchesani syndrome. Hum Mol Genet. 2018;27(21):3675–87. Epub 2018/07/31. doi: 10.1093/hmg/ddy276. PubMed PMID: 30060141; PMCID: PMC6196651.

30. Wang LW, Kutz WE, Mead TJ, Beene LC, Singh S, Jenkins MW, Reinhardt DP, Apte SS. Adamts10 inactivation in mice leads to persistence of ocular microfibrils subsequent to reduced fibrillin-2 cleavage. Matrix Biol. 2019;77:117–28. Epub 2018/09/12. doi: 10.1016/j.matbio.2018.09.004. PubMed PMID: 30201140.

31. Prins BP, Mead TJ, Brody JA, Sveinbjornsson G, Ntalla I, Bihlmeyer NA, van den Berg M, Bork-Jensen J, Cappellani S, Van Duijvenboden S, Klena NT, Gabriel GC, Liu X, Gulec C, Grarup N, Haessler J, Hall LM, Iorio A, Isaacs A, Li-Gao R, Lin H, Liu CT, Lyytikäinen LP, Marten J, Mei H, Müller-Nurasyid M, Orini M, Padmanabhan S, Radmanesh F, Ramirez J, Robino A, Schwartz M, van Setten J, Smith AV, Verweij N, Warren HR, Weiss S, Alonso A, Arnar DO, Bots ML, de Boer RA, Dominiczak AF, Eijgelsheim M, Ellinor PT, Guo X, Felix SB, Harris TB, Hayward C, Heckbert SR, Huang PL, Jukema JW, Kähönen M, Kors JA, Lambiase PD, Launer LJ, Li M, Linneberg A, Nelson CP, Pedersen O, Perez M, Peters A, Polasek O, Psaty BM, Raitakari OT, Rice KM, Rotter JI, Sinner MF, Soliman EZ, Spector TD, Strauch K, Thorsteinsdottir U, Tinker A, Trompet S, Uitterlinden A, Vaartjes I, van der Meer P, Völker U, Völzke H, Waldenberger M, Wilson JG, Xie Z, Asselbergs FW, Dörr M, van Duijn CM, Gasparini P, Gudbjartsson DF, Gudnason V, Hansen T, Kääb S, Kanters JK, Kooperberg C, Lehtimäki T, Lin HJ, Lubitz SA, Mook-Kanamori DO, Conti FJ, Newton-Cheh CH, Rosand J, Rudan I, Samani NJ, Sinagra G, Smith BH, Holm H, Stricker BH, Ulivi S, Sotoodehnia N, Apte SS, van der Harst P, Stefansson K, Munroe PB, Arking DE, Lo CW, Jamshidi Y. Exome-chip meta-analysis identifies novel loci associated with cardiac conduction, including ADAMTS6. Genome Biol. 2018;19(1):87. Epub 2018/07/18. doi: 10.1186/s13059-018-1457-6. PubMed PMID: 30012220; PMCID: PMC6048820.

32. Dubail J, Aramaki-Hattori N, Bader HL, Nelson CM, Katebi N, Matuska B, Olsen BR, Apte SS. A new Adamts9 conditional mouse allele identifies its non-redundant role in interdigital web regression. Genesis. 2014;52(7):702–12. doi: 10.1002/dvg.22784. PubMed PMID: 24753090; PMCID: 4107014.

33. Enomoto H, Nelson, C., Somerville, R.P.T., Mielke, K., Dixon, L., Powell, K., Apte, S.S.. Cooperation of two ADAMTS metalloproteases in closure of the mouse palate identifies a requirement for versican proteolysis in regulating palatal mesenchyme proliferation. Development. 2010;137:4029–38.

34. McCulloch DR, Nelson CM, Dixon LJ, Silver DL, Wylie JD, Lindner V, Sasaki T, Cooley MA, Argraves WS, Apte SS. ADAMTS metalloproteases generate active versican fragments that regulate interdigital web regression. Dev Cell. 2009;17(5):687–98. Epub 2009/11/20. doi: S1534-5807(09)00390-6 [pii] 10.1016/j.devcel.2009.09.008. PubMed PMID: 19922873; PMCID: 2780442.

35. Mead TJ, McCulloch DR, Ho JC, Du Y, Adams SM, Birk DE, Apte SS. The metalloproteinase-proteoglycans ADAMTS7 and ADAMTS12 provide an innate, tendon-specific protective mechanism against heterotopic ossification. JCI Insight. 2018;3(7). Epub 2018/04/06. doi: 10.1172/jci.insight.92941. PubMed PMID: 29618652; PMCID: PMC5928868.

36. Nandadasa S, Kraft CM, Wang LW, O’Donnell A, Patel R, Gee HY, Grobe K, Cox TC, Hildebrandt F, Apte SS. Secreted metalloproteases ADAMTS9 and ADAMTS20 have a non-canonical role in ciliary vesicle growth during ciliogenesis. Nat Commun. 2019;10(1):953. Epub 2019/03/01. doi: 10.1038/s41467-019-08520-7. PubMed PMID: 30814516; PMCID: PMC6393521.

37. Nandadasa S, Nelson CM, Apte SS. ADAMTS9-Mediated Extracellular Matrix Dynamics Regulates Umbilical Cord Vascular Smooth Muscle Differentiation and Rotation. Cell Rep. 2015;11(10):1519–28. doi: 10.1016/j.celrep.2015.05.005. PubMed PMID: 26027930; PMCID: PMC4472575.

38. El-Brolosy MA, Kontarakis Z, Rossi A, Kuenne C, Günther S, Fukuda N, Kikhi K, Boezio GLM, Takacs CM, Lai SL, Fukuda R, Gerri C, Giraldez AJ, Stainier DYR. Genetic compensation triggered by mutant mRNA degradation. Nature. 2019;568(7751):193–7. Epub 2019/04/05. doi: 10.1038/s41586-019-1064-z. PubMed PMID: 30944477; PMCID: PMC6707827.

39. Sztal TE, Stainier DYR. Transcriptional adaptation: a mechanism underlying genetic robustness. Development. 2020;147(15). Epub 2020/08/21. doi: 10.1242/dev.186452. PubMed PMID: 32816903.

40. Somerville RP, Jungers KA, Apte SS. ADAMTS10: Discovery and characterization of a novel, widely expressed metalloprotease and its proteolytic activation. J Biol Chem. 2004;279:51208–17. PubMed PMID: 15355968.

41. Mularczyk EJ, Singh M, Godwin ARF, Galli F, Humphreys N, Adamson AD, Mironov A, Cain SA, Sengle G, Boot-Handford RP, Cossu G, Kielty CM, Baldock C. ADAMTS10-mediated tissue disruption in Weill-Marchesani Syndrome. Hum Mol Genet. 2018. doi: 10.1093/hmg/ddy276. PubMed PMID: 30060141.

42. Jensen SA, Reinhardt DP, Gibson MA, Weiss AS. Protein interaction studies of MAGP-1 with tropoelastin and fibrillin-1. J Biol Chem. 2001;276(43):39661–6. Epub 2001/08/02. doi: 10.1074/jbc.M104533200 M104533200 [pii]. PubMed PMID: 11481325.

43. Beene LC, Wang LW, Hubmacher D, Keene DR, Reinhardt DP, Annis DS, Mosher DF, Mecham RP, Traboulsi EI, Apte SS. Nonselective assembly of fibrillin 1 and fibrillin 2 in the rodent ocular zonule and in cultured cells: implications for marfan syndrome. Invest Ophthalmol Vis Sci. 2013;54(13):8337–44. doi: 10.1167/iovs.13-13121. PubMed PMID: 24265020; PMCID: 3875392.

44. Sabatier L, Chen D, Fagotto-Kaufmann C, Hubmacher D, McKee MD, Annis DS, Mosher DF, Reinhardt DP. Fibrillin assembly requires fibronectin. Mol Biol Cell. 2009;20(3):846–58. Epub 2008/11/28. doi: E08-08-0830 [pii] 10.1091/mbc.E08-08-0830. PubMed PMID: 19037100; PMCID: 2633374.

45. Cain SA, Morgan A, Sherratt MJ, Ball SG, Shuttleworth CA, Kielty CM. Proteomic analysis of fibrillin-rich microfibrils. Proteomics. 2006;6(1):111–22. Epub 2005/11/23. doi: 10.1002/pmic.200401340. PubMed PMID: 16302274.

46. De Maria A, Wilmarth PA, David LL, Bassnett S. Proteomic Analysis of the Bovine and Human Ciliary Zonule. Invest Ophthalmol Vis Sci. 2017;58(1):573–85. doi: 10.1167/iovs.16-20866. PubMed PMID: 28125844; PMCID: PMC5283081.

47. Fujikawa Y, Yoshida H, Inoue T, Ohbayashi T, Noda K, von Melchner H, Iwasaka T, Shiojima I, Akama TO, Nakamura T. Latent TGF-β binding protein 2 and 4 have essential overlapping functions in microfibril development. SciRep. 2017;7:43714. Epub 2017/03/03. doi: 10.1038/srep43714. PubMed PMID: 28252045; PMCID: PMC5333096.

48. Mecham RP, Gibson MA. The microfibril-associated glycoproteins (MAGPs) and the microfibrillar niche. Matrix Biol. 2015;47:13–33. Epub 2015/05/13. doi: 10.1016/j.matbio.2015.05.003. PubMed PMID: 25963142; PMCID: PMC5154765.

49. Kuno K, Matsushima K. ADAMTS-1 protein anchors at the extracellular matrix through the thrombospondin type I motifs and its spacing region. J Biol Chem. 1998;273(22):13912–7.

50. Kleifeld O, Doucet A, auf dem Keller U, Prudova A, Schilling O, Kainthan RK, Starr AE, Foster LJ, Kizhakkedathu JN, Overall CM. Isotopic labeling of terminal amines in complex samples identifies protein N-termini and protease cleavage products. Nat Biotechnol. 2010;28(3):281–8. doi: 10.1038/nbt.1611. PubMed PMID: 20208520.

51. Kockmann T, Carte, N., Melkko, S. & Keller, U.a.d.. Identification of Protease Substrates in Complex Proteomes by iTRAQ-TAILs on a Thermo Q Exactive Instrument In: Grant JE, Li, H., editor. Neuromethods New York: Springer Science+Business Media; 2016. p. 187–207.

52. Bekhouche M, Leduc C, Dupont L, Janssen L, Delolme F, Vadon-Le Goff S, Smargiasso N, Baiwir D, Mazzucchelli G, Zanella-Cleon I, Dubail J, De Pauw E, Nusgens B, Hulmes DJ, Moali C, Colige A. Determination of the substrate repertoire of ADAMTS2, 3, and 14 significantly broadens their functions and identifies extracellular matrix organization and TGF-beta signaling as primary targets. FASEB J. 2016;30(5):1741-56. doi: 10.1096/fj.15-279869. PubMed PMID: 26740262.

53. Hubmacher D, El-Hallous EI, Nelea V, Kaartinen MT, Lee ER, Reinhardt DP. Biogenesis of extracellular microfibrils: Multimerization of the fibrillin-1 C terminus into bead-like structures enables self-assembly. Proc Natl Acad Sci U S A. 2008;105(18):6548–53. Epub 2008/05/02. doi: 0706335105 [pii] 10.1073/pnas.0706335105. PubMed PMID: 18448684; PMCID: 2373353.

54. Chun TH, Sabeh F, Ota I, Murphy H, McDonagh KT, Holmbeck K, Birkedal-Hansen H, Allen ED, Weiss SJ. MT1-MMP-dependent neovessel formation within the confines of the three-dimensional extracellular matrix. J Cell Biol. 2004;167(4):757–67. Epub 2004/11/17. doi: 10.1083/jcb.200405001. PubMed PMID: 15545316; PMCID: PMC2172577.

55. Inada M, Wang Y, Byrne MH, Rahman MU, Miyaura C, Lopez-Otin C, Krane SM. Critical roles for collagenase-3 (Mmp13) in development of growth plate cartilage and in endochondral ossification. Proc Natl Acad Sci U S A. 2004;101(49):17192–7. PubMed PMID: 15563592.

56. Nandadasa S, Foulcer S, Apte SS. The multiple, complex roles of versican and its proteolytic turnover by ADAMTS proteases during embryogenesis. Matrix Biol. 2014;35:34–41. doi: 10.1016/j.matbio.2014.01.005. PubMed PMID: 24444773.

57. Cain SA, Baldock C, Gallagher J, Morgan A, Bax DV, Weiss AS, Shuttleworth CA, Kielty CM. Fibrillin-1 interactions with heparin. Implications for microfibril and elastic fiber assembly. J Biol Chem. 2005;280(34):30526–37. Epub 2005/06/28. doi: M501390200 [pii] 10.1074/jbc.M501390200. PubMed PMID: 15980072.

58. Dallas SL, Keene DR, Bruder SP, Saharinen J, Sakai LY, Mundy GR, Bonewald LF. Role of the latent transforming growth factor beta binding protein 1 in fibrillin-containing microfibrils in bone cells in vitro and in vivo. J Bone Miner Res. 2000;15(1):68–81. Epub 2000/01/26. PubMed PMID: 10646116.

59. Kettle S, Yuan X, Grundy G, Knott V, Downing AK, Handford PA. Defective calcium binding to fibrillin-1: consequence of an N2144S change for fibrillin-1 structure and function. J Mol Biol. 1999;285(3):1277–87. Epub 1999/01/15. doi: S0022-2836(98)92368-3 [pii] 10.1006/jmbi.1998.2368. PubMed PMID: 9887276.

60. Kinsey R, Williamson MR, Chaudhry S, Mellody KT, McGovern A, Takahashi S, Shuttleworth CA, Kielty CM. Fibrillin-1 microfibril deposition is dependent on fibronectin assembly. J Cell Sci. 2008;121(Pt 16):2696–704. Epub 2008/07/26. doi: jcs.029819 [pii] 10.1242/jcs.029819. PubMed PMID: 18653538.

61. Charbonneau NL, Jordan CD, Keene DR, Lee-Arteaga S, Dietz HC, Rifkin DB, Ramirez F, Sakai LY. Microfibril structure masks fibrillin-2 in postnatal tissues. J Biol Chem. 2010;285(26):20242–51. Epub 2010/04/21. doi: 10.1074/jbc.M109.087031. PubMed PMID: 20404337; PMCID: PMC2888437.

62. Faivre L, Dollfus H, Lyonnet S, Alembik Y, Megarbane A, Samples J, Gorlin RJ, Alswaid A, Feingold J, Le Merrer M, Munnich A, Cormier-Daire V. Clinical homogeneity and genetic heterogeneity in Weill-Marchesani syndrome. Am J Med Genet. 2003;123A(2):204–7. PubMed PMID: 14598350.

63. Bader HL, Ruhe AL, Wang LW, Wong AK, Walsh KF, Packer RA, Mitelman J, Robertson KR, O’Brien DP, Broman KW, Shelton GD, Apte SS, Neff MW. An ADAMTSL2 founder mutation causes Musladin-Lueke Syndrome, a heritable disorder of beagle dogs, featuring stiff skin and joint contractures. PLoS One. 2010;5(9): e12817. Epub 2010/09/24. doi: 10.1371/journal.pone.0012817. PubMed PMID: 20862248; PMCID: 2941456.

64. Bader HL, Wang LW, Ho JC, Tran T, Holden P, Fitzgerald J, Atit RP, Reinhardt DP, Apte SS. A disintegrin-like and metalloprotease domain containing thrombospondin type 1 motif-like 5 (ADAMTSL5) is a novel fibrillin-1-, fibrillin-2-, and heparin-binding member of the ADAMTS superfamily containing a netrin-like module. Matrix Biol. 2012;31(7-8):398–411. doi: 10.1016/j.matbio.2012.09.003. PubMed PMID: 23010571; PMCID: 3546522.

65. Gabriel LA, Wang LW, Bader H, Ho JC, Majors AK, Hollyfield JG, Traboulsi EI, Apte SS. ADAMTSL4, a secreted glycoprotein widely distributed in the eye, binds fibrillin-1 microfibrils and accelerates microfibril biogenesis. Invest Ophthalmol Vis Sci. 2012;53(1):461–9. Epub 2011/10/13. doi: iovs.10-5955 [pii] 10.1167/iovs.10-5955. PubMed PMID: 21989719.

66. Le Goff C, Morice-Picard F, Dagoneau N, Wang LW, Perrot C, Crow YJ, Bauer F, Flori E, Prost-Squarcioni C, Krakow D, Ge G, Greenspan DS, Bonnet D, Le Merrer M, Munnich A, Apte SS, Cormier-Daire V. ADAMTSL2 mutations in geleophysic dysplasia demonstrate a role for ADAMTS-like proteins in TGF-beta bioavailability regulation. Nat Genet. 2008;40(9):1119–23. Epub 2008/08/05. doi: 10.1038/ng.199. PubMed PMID: 18677313; PMCID: 2675613.

67. Saito M, Kurokawa M, Oda M, Oshima M, Tsutsui K, Kosaka K, Nakao K, Ogawa M, Manabe R, Suda N, Ganjargal G, Hada Y, Noguchi T, Teranaka T, Sekiguchi K, Yoneda T, Tsuji T. ADAMTSL6beta protein rescues fibrillin-1 microfibril disorder in a Marfan syndrome mouse model through the promotion of fibrillin-1 assembly. J Biol Chem. 2011;286(44):38602–13. Epub 2011/09/02. doi: M111.243451 [pii] 10.1074/jbc.M111.243451. PubMed PMID: 21880733; PMCID: 3207443.

68. Tsutsui K, Manabe RI, Yamada T, Nakano I, Oguri Y, Keene DR, Sengle G, Sakai LY, Sekiguchi K. A disintegrin and metalloproteinase with thrombospondin motifs-like-6 (ADAMTSL-6) is a novel extracellular matrix protein that binds to fibrillin-1 and promotes fibrillin-1 fibril formation. J Biol Chem. 2010;285:4870–82. Epub 2009/11/27. doi: M109.076919 [pii] 10.1074/jbc.M109.076919. PubMed PMID: 19940141.

69. Mead TJ, McCulloch DR, Ho JC, Du Y, Adams SM, Birk DE, Apte SS. The metalloproteinase-proteoglycans ADAMTS7 and ADAMTS12 provide an innate, tendon-specific protective mechanism against heterotopic ossification. JCI Insight. 2018;3(7):e92941. doi: 10.1172/jci.insight.92941. PubMed PMID: 29618652.

70. Gregory KE, Ono RN, Charbonneau NL, Kuo CL, Keene DR, Bachinger HP, Sakai LY. The prodomain of BMP-7 targets the BMP-7 complex to the extracellular matrix. J Biol Chem. 2005;280(30):27970–80. PubMed PMID: 15929982.

71. Sengle G, Ono RN, Sasaki T, Sakai LY. Prodomains of transforming growth factor beta (TGFbeta) superfamily members specify different functions: extracellular matrix interactions and growth factor bioavailability. J Biol Chem. 2011;286(7):5087–99. Epub 2010/12/08. doi: M110.188615 [pii] 10.1074/jbc.M110.188615. PubMed PMID: 21135108; PMCID: 3037620.

72. Furlan AG, Spanou CES, Godwin ARF, Wohl AP, Zimmermann LA, Imhof T, Koch M, Baldock C, Sengle G. A new MMP-mediated prodomain cleavage mechanism to activate bone morphogenetic proteins from the extracellular matrix. Faseb j. 2021;35(3):e21353. Epub 2021/02/26. doi: 10.1096/fj.202001264R. PubMed PMID: 33629769.

73. Wohl AP, Troilo H, Collins RF, Baldock C, Sengle G. Extracellular Regulation of Bone Morphogenetic Protein Activity by the Microfibril Component Fibrillin-1. J Biol Chem. 2016;291(24):12732–46. Epub 2016/04/10. doi: 10.1074/jbc.M115.704734. PubMed PMID: 27059954; PMCID: PMC4933460.

74. Colnot C, de la Fuente L, Huang S, Hu D, Lu C, St-Jacques B, Helms JA. Indian hedgehog synchronizes skeletal angiogenesis and perichondrial maturation with cartilage development. Development. 2005;132(5):1057–67. Epub 2005/02/04. doi: 10.1242/dev.01649. PubMed PMID: 15689378.

75. Maes C. Signaling pathways effecting crosstalk between cartilage and adjacent tissues: Seminars in cell and developmental biology: The biology and pathology of cartilage. Semin Cell Dev Biol. 2017;62:16–33. Epub 2016/05/18. doi: 10.1016/j.semcdb.2016.05.007. PubMed PMID: 27180955.

76. Vortkamp A, Lee K, Lanske B, Segre GV, Kronenberg HM, Tabin CJ. Regulation of rate of cartilage differentiation by Indian hedgehog and PTH-related protein. Science. 1996;273(5275):613-22. Epub 1996/08/02. doi: 10.1126/science.273.5275.613. PubMed PMID: 8662546.

77. Murphy MK, Huey DJ, Hu JC, Athanasiou KA. TGF-β1, GDF-5, and BMP-2 stimulation induces chondrogenesis in expanded human articular c^β^hondrocytes and marrow-derived stromal cells. Stem Cells. 2015;33(3):762–73. Epub 2014/11/08. doi: 10.1002/stem.1890. PubMed PMID: 25377511.

78. Hatakeyama Y, Tuan RS, Shum L. Distinct functions of BMP4 and GDF5 in the regulation of chondrogenesis. J Cell Biochem. 2004;91(6):1204–17. Epub 2004/03/30. doi: 10.1002/jcb.20019. PubMed PMID: 15048875.

79. Miyamoto A, Lau R, Hein PW, Shipley JM, Weinmaster G. Microfibrillar proteins MAGP-1 and MAGP-2 induce Notch1 extracellular domain dissociation and receptor activation. J Biol Chem. 2006;281(15):10089–97. doi: 10.1074/jbc.M600298200. PubMed PMID: 16492672.

80. Nehring LC, Miyamoto A, Hein PW, Weinmaster G, Shipley JM. The extracellular matrix protein MAGP-2 interacts with Jagged1 and induces its shedding from the cell surface. J Biol Chem. 2005;280(21):20349–55. doi: 10.1074/jbc.M500273200. PubMed PMID: 15788413.

81. Pereira L, Lee SY, Gayraud B, Andrikopoulos K, Shapiro SD, Bunton T, Biery NJ, Dietz HC, Sakai LY, Ramirez F. Pathogenetic sequence for aneurysm revealed in mice underexpressing fibrillin-1. Proc Natl Acad Sci U S A. 1999;96(7):3819–23. PubMed PMID: 10097121.

82. Li Y, Klena NT, Gabriel GC, Liu X, Kim AJ, Lemke K, Chen Y, Chatterjee B, Devine W, Damerla RR, Chang C, Yagi H, San Agustin JT, Thahir M, Anderton S, Lawhead C, Vescovi A, Pratt H, Morgan J, Haynes L, Smith CL, Eppig JT, Reinholdt L, Francis R, Leatherbury L, Ganapathiraju MK, Tobita K, Pazour GJ, Lo CW. Global genetic analysis in mice unveils central role for cilia in congenital heart disease. Nature. 2015;521(7553):520–4. doi: 10.1038/nature14269. PubMed PMID: 25807483; PMCID: PMC4617540.

83. Gaytan F, Morales C, Reymundo C, Tena-Sempere M. A novel RGB-trichrome staining method for routine histological analysis of musculoskeletal tissues. Sci Rep. 2020;10(1):16659. Epub 2020/10/09. doi: 10.1038/s41598-020-74031-x. PubMed PMID: 33028938; PMCID: PMC7541469.

84. Mead TJ, Yutzey KE. Notch pathway regulation of chondrocyte differentiation and proliferation during appendicular and axial skeleton development. Proc Natl Acad Sci U S A. 2009;106(34):14420–5. doi: 10.1073/pnas.0902306106. PubMed PMID: 19590010; PMCID: 2732875.

85. Schindelin J, Arganda-Carreras I, Frise E, Kaynig V, Longair M, Pietzsch T, Preibisch S, Rueden C, Saalfeld S, Schmid B, Tinevez JY, White DJ, Hartenstein V, Eliceiri K, Tomancak P, Cardona A. Fiji: an open-source platform for biological-image analysis. Nat Methods. 2012;9(7):676–82. Epub 2012/06/30. doi: 10.1038/nmeth.2019. PubMed PMID: 22743772; PMCID: PMC3855844.

86. Wang W, Chun H, Baek J, Sadik JE, Shirazyan A, Razavi P, Lopez N, Lyons KM. The TGFβ type I receptor TGFβRI functions as an inhibitor of BMP signaling in cartilage. Proc Natl Acad Sci U S A. 2019;116(31):15570–9. Epub 2019/07/18. doi: 10.1073/pnas.1902927116. PubMed PMID: 31311865; PMCID: PMC6681752.

87. Mead TJ. Alizarin Red and Alcian Blue Preparations to Visualize the Skeleton. Methods Mol Biol. 2020;2043:207–12. Epub 2019/08/30. doi: 10.1007/978-1-4939-9698-8_17. PubMed PMID: 31463914.

88. Mead TJ, Du Y, Nelson CM, Gueye NA, Drazba J, Dancevic CM, Vankemmelbeke M, Buttle DJ, Apte SS. ADAMTS9-Regulated Pericellular Matrix Dynamics Governs Focal Adhesion-Dependent Smooth Muscle Differentiation. Cell Rep. 2018;23(2):485–98. doi: 10.1016/j.celrep.2018.03.034. PubMed PMID: 29642006.

89. Mead TJ, Apte SS. Expression Analysis by RNAscope In Situ Hybridization. Methods Mol Biol. 2020;2043:173–8. Epub 2019/08/30. doi: 10.1007/978-1-4939-9698-8_14. PubMed PMID: 31463911.

90. Hubmacher D, Schneider M, Berardinelli SJ, Takeuchi H, Willard B, Reinhardt DP, Haltiwanger RS, Apte SS. Unusual life cycle and impact on microfibril assembly of ADAMTS17, a secreted metalloprotease mutated in genetic eye disease. Sci Rep. 2017;7:41871. doi: 10.1038/srep41871. PubMed PMID: 28176809; PMCID: PMC5296908.

91. Wang LW, Nandadasa S, Annis DS, Dubail J, Mosher DF, Willard BB, Apte SS. A disintegrin-like and metalloproteinase domain with thrombospondin type 1 motif 9 (ADAMTS9) regulates fibronectin fibrillogenesis and turnover. J Biol Chem. 2019;294(25):9924–36. Epub 2019/05/16. doi: 10.1074/jbc.RA118.006479. PubMed PMID: 31085586; PMCID: PMC6597835.

92. Martin DR, Witten JC, Tan CD, Rodriguez ER, Blackstone EH, Pettersson GB, Seifert DE, Willard BB, Apte SS. Proteomics identifies a convergent innate response to infective endocarditis and extensive proteolysis in vegetation components. JCI Insight. 2020;5(14). Epub 2020/06/17. doi: 10.1172/jci.insight.135317. PubMed PMID: 32544089; PMCID: PMC7453909.

93. Kockmann T, Carte N, Melkko S, auf dem Keller U. Identification of Protease Substrates in Complex Proteomes by iTRAQ-TAILS on a Thermo Q Exactive Instrument. In: Grant JE, Li H, editors. Analysis of Post-Translational Modifications and Proteolysis in Neuroscience. New York, NY: Springer New York; 2016. p. 187–207.

94. Trask TM, Trask BC, Ritty TM, Abrams WR, Rosenbloom J, Mecham RP. Interaction of tropoelastin with the amino-terminal domains of fibrillin-1 and fibrillin-2 suggests a role for the fibrillins in elastic fiber assembly. J Biol Chem. 2000;275(32):24400–6. Epub 2000/05/29. doi: 10.1074/jbc.M003665200. PubMed PMID: 10825173.

95. Shi Y, Jones W, Beatty W, Tan Q, Mecham RP, Kumra H, Reinhardt DP, Gibson MA, Reilly MA, Rodriguez J, Bassnett S. Latent-transforming growth factor beta-binding protein-2 (LTBP-2) is required for longevity but not for development of zonular fibers. Matrix Biol. 2020. Epub 2020/10/12. doi: 10.1016/j.matbio.2020.10.002. PubMed PMID: 33039488.

96. Trask BC, Trask TM, Broekelmann T, Mecham RP. The microfibrillar proteins MAGP-1 and fibrillin-1 form a ternary complex with the chondroitin sulfate proteoglycan decorin. Mol Biol Cell. 2000;11(5):1499–507. Epub 2000/05/04. doi: 10.1091/mbc.11.5.1499. PubMed PMID: 10793130; PMCID: PMC14862.

